# GISA: Using Gauss Integrals to identify rare conformations in protein structures

**DOI:** 10.1101/758029

**Authors:** Christian Grønbæk, Thomas Hamelryck, Peter Røgen

## Abstract

The native structure of a protein is important for its function, and therefore methods for exploring protein structures have attracted much research. However, rather few methods are sensitive to topologic-geometric features, the examples being knots, slipknots, lassos, links, and pokes, and with each method aimed only for a specific set of such configurations.

We here propose a general method which transforms a structure into a “fingerprint of topological-geometric values” consisting in a series of real-valued descriptors from mathematical Knot Theory. The extent to which a structure contains unusual configurations can then be judged from this fingerprint. The method is therefore not confined to a particular pre-defined topology or geometry (like a knot or a poke), and so, unlike existing methods, it is general. To achieve this our new algorithm, GISA, as a key novelty produces the descriptors, so called Gauss integrals, not only for the full chains of a protein but for all its sub-chains, thereby allowing fingerprinting on any scale from local to global. The Gauss integrals are known to be effective descriptors of global protein folds.

Applying GISA to a set of about 8000 high resolution structures (top8000), we first show how it enables swift identification of predefined geometries such as pokes and links. We then apply GISA with no restrictions on geometry, to show how it allows identifying rare conformations by finding rare invariant values only. In this unrestricted search, pokes and links are still found, but also knotted conformations, as well as more highly entangled configurations not previously described. Thus, applying the basic scan method in GISA’s tool-box to the top8000 set, 10 known cases of knots are ranked as the top positive Gauss number cases, while placing at the top of the negative Gauss numbers 14 cases in cis-trans isomerases sharing a spatial motif of little secondary structure content, which possibly has gone unnoticed.

Potential applications of the GISA tools include finding errors in protein models and identifying unusual conformations that might be important for protein folding and function. By its broad potential, we believe that GISA will be of general benefit to the structural bioinformatics community.

GISA is coded in C and comes as a command line tool. Source and compiled code for GISA plus read-me and examples are publicly available at GitHub (https://github.com).

## Background

In [12] P.Røgen and H.Bohr introduced a set of quantitative protein fold descriptors consisting in 29 knot-theoretic Gauss Integral (GI) based invariants, shown shortly after in [7] to be able to automatically recover the classification of the CATH database.

Automated *local* scrutiny of folds is desired for various purposes, including identifying odd shapes in predictions, improving classification and for structure alignments. However, while the GI invariants work very well as *global* fold descriptors an efficient method for computing them locally has been lacking and local applications have been few. Revealing though indeed the relevance of applying GIs locally, the structural similarity methods in [1] and [11] based on local writhe showed excellent performance (writhe is an order one GI).

Due to its recursive nature, our new algorithm, GISA, computes not only the GI invariants of an entire chain but at the same time of all sub-chains, allowing therefore structural analyses on any scale from local to global (we assume chains and sub-chains to be connected).

By the knot-theoretic nature of the GIs, GISA is sensitive to topologic-geometric differences, while having the fundamental translational-rotational invariance. This distinguishes GISA from distance based approaches, and comparisons e.g. to methods building on RMSD are therefore not meaningful. A general method for structural analysis having such topologic-geometric sensitivity seems still to be lacking ([3] [6]). By its versatility, we believe that GISA can fill this gap.

As a tool GISA includes computation of the desired GIs, deriving GI-values for sub-chains and search methods allowing to rank a set of query structures’ GI-values against a back ground, consisting in a set of GI-values, likewise produced by running GISA on a “data base” of structures.

The main aims of this paper are to introduce GISA, to explain the method via a proof-of-concept and describe the contents of the tool. The main focus is on the proof-of-concept test: First we show in a “restricted search” how GISA can be exploited to provide an algorithm for identifying particular geometries in folds such as the “pokes” and “co-pokes” pointed out by F.Khatib et al. [2]. Then, in a similar unrestricted search, we show how GISA allows identifying the very same configurations as well as more elaborate ones, by letting the search being based on finding outlying writhe values only (see Figure 1 and Figure 2 below; more examples from both searches can be found in the Supporting Material).

**Figure 1:**
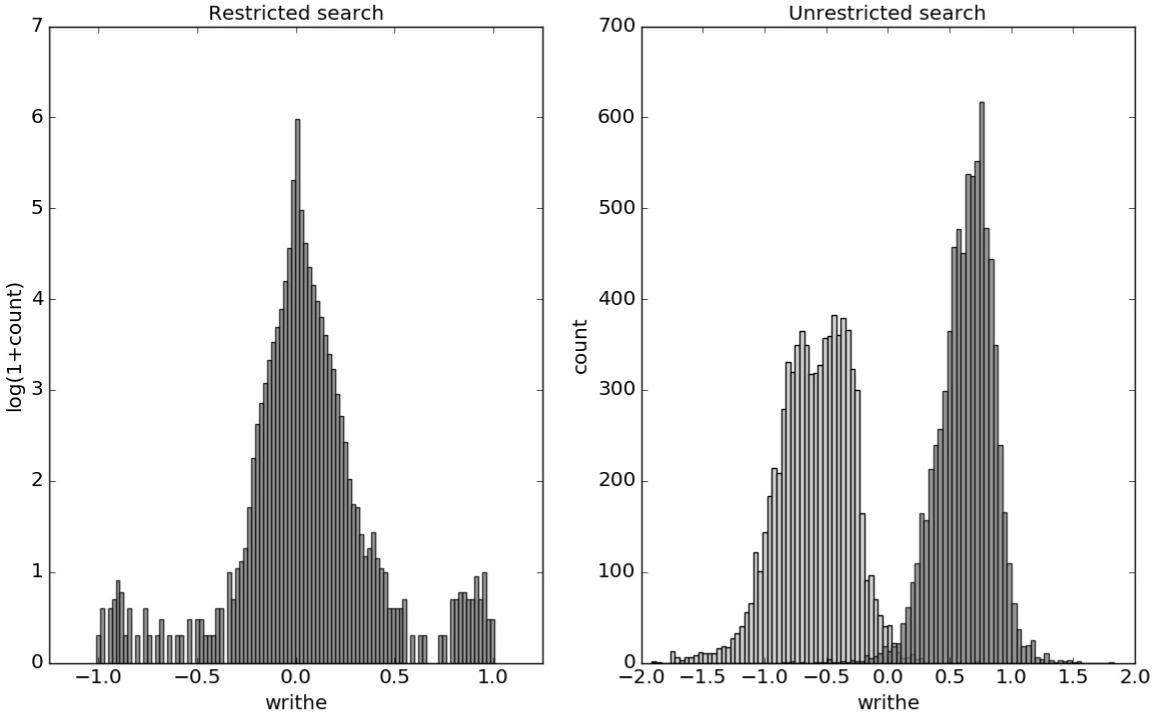
Distributions of mutual writhe values for potential links (left, restricted search) and for pairs of sub-chains of length 30 (right, unrestricted search) throughout the top8000 set, obtained as described in the text. For the unrestricted search the light-grey (dark-grey) bars show the distribution of the lowest (highest) writhe value per chain (see text).

**Figure 2:**
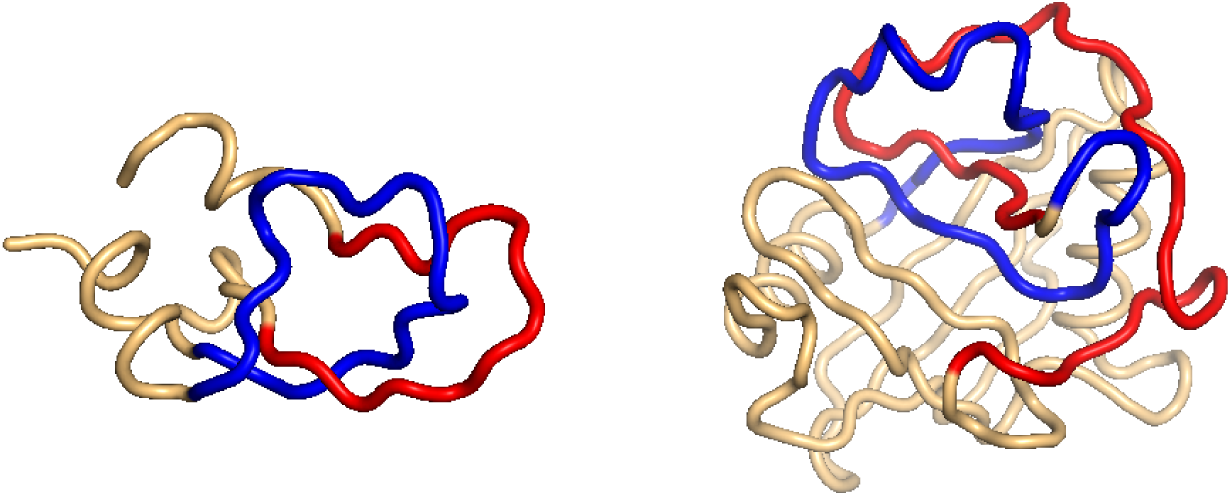
Two examples of particularly high mutual writhe value between the red and the blue sub-chains. Left: a 1-link in the rather small 1bpi structure. Right: an example of more elaborate geometry in the 2cpl protein found using the unrestricted search method. The plots were made with Pymol [13].

As a natural continuation of the proof-of-concept, GISA’s basic scan tool rar0 (rar is short for rarity) formalizes the unrestricted search so as to allow assessing a set of structures on a larger back ground by means of the value of the order one GI only (the writhe). The other main tools in GISA are the GI-generating functions and two more advanced scan methods (rar1 and rar2). All three scan tools build on the possibility of pre-computing the GIs so as to generate a back ground against which query sets can be swiftly assessed. Key in the pre-computatibility of the GIs is their translational and rotational invariance (no costly superpositions of the structures are needed). While our focus in this paper is on the application of the writhe, and hence rar0, the two more advanced scan flavours allow including higher order GIs. These both build on discretizing the GI-arrays and looking up the resulting GI-words in the likewise “dictionaried” back ground of GI-arrays. This can be seen as local structural alignment in the space of GI-arrays (rather than in 3d-space) and could serve as the foundation of a similar global structural alignment. As auxiliaries GISA includes tools to support the two search types involved in the proof-of-concept.

To validate the output of the restricted and unrestricted searches we visually inspect the cases having the most conspicuous GI values and compare their rankings. To assess GISA’s scan methods, we show that their outputs are well-aligned at similar settings, and in particular that the top-rankings of the basic scan tool rar0 match those of the unrestricted search.

Recent work by P.Dabrowski-Tumanski and J.I.Sulkowska [8] indicates the steady interest in links and closely related topologies. While the search algorithms in [2] and [8] build on predefined shapes, our algorithm searches for exceptional values of GIs, and the shapes of the search hits are then final output rather than input; thus GISA allows to identify both less constrained as well as more elaborate shapes. This we support by running the unrestricted search and, more generally, by the runs of GISA’s in-built scan methods.

The command line tools (compiled code) as well as the source C code for computing the invariants up to and including order 3, for supporting the scans and for the particular searches are available at GitHub under GNU General Public License v3 (https://github.com).

## Methods

### Approach in restricted and unrestricted searches

We represent a protein chain by the piece-wise linear curve given by its α-Carbon trace. A poke consists of an almost closed loop (of moderate length) through which a shorter segment of the chain sticks — or pokes. A co-poke can be understood as a “1-link”: two almost closed loops (of moderate length) poking through each other, i.e. interlinking once. We prefer the term 1-link for the following reason.

The relevance of the Gauss Integrals is that in a 1-link the two smooth closed loops have a linking number, or writhe, of *±*1. If the loops are only almost closed, the writhe will be close to *±*1, and, loosely speaking, the writhe increases (in absolute value) the more the two curves interlink. Therefore, if one can compute the writhe values of all pairs of (almost) closed loops in a chain, instances of (almost) 1-links can possibly be identified (cases of higher linking numbers — i.e. one loop winding around the other several times — are less expected due to their lower entropy). A major benefit of our GISA algorithm is that it allows exactly that: it computes not only Gauss Integrals of the whole chain, but of all sub-chains. The governing equation of the algorithm (Supporting Material, eq.1, p.4) then allows to swiftly compute the desired *mutual* writhe value of any given pair of sub-chains.

Pokes can be searched for similarly by considering the mutual writhe values of short sub-chains (length 10, say) vs. (almost) closed loops. The intuition here is that of the electromagnetic induction in a piece of wire poking through a loop of another wire through which an electrical current runs; the “purer” the poke, the higher the induction.

The base of GISA consists in an algorithm computing the GIs, which includes also support for the two modes: 1) for the restricted search, all (almost) closed loops are identified, whereafter the relevant mutual writhe values for link/poke searching are derived and written to file; 2) for the unrestricted search, it simply derives the mutual writhe of all pairs of sub-chains of some fixed length. To limit the size of the output, we keep only the pairs of sub-chains with the lowest and with the highest mutual writhe values. The derivation of the mutual writhe amounts to adding four look-ups in the stored writhe-table. In both versions GISA returns the identified sub-chain pairs along with their mutual writhe. Conspicuous cases are then found in the tails of the resulting distribution of mutual writhe values. In the restricted search we expect to find 1-links/pokes and, when using the unrestricted search, possibly other geometries as well.

While a small downstream task is here needed to identify tail events, GISA’s scan methods that we now turn to directly rank structures in terms of their “GI content”. Of these the basic method (rar0) is the offspring of our proof-of-concept.

### GISA’s scan methods

As explained GISA includes tools for querying possible rarity in a set of structures against a preferably larger and representative set of structures. The output is a score and a probability/rank for each queried structure, with all queried structures listed by decreasing score (and increasing probability/rank). GISA has three types of scans to support this, rar0, rar1 and rar2 (in the command line tool these are referred to as flavours). In all three, sub-chains (windows) of a set length are used, covering each structure by moving the window along the structure at a likewise pre-set step size.

The basic scan method (rar0) assesses each structure by its content of mutual writhe above a set threshold. There are two versions. In one (A) we distinguish between pairs of positive and negative mutual writhe. Each structure is here represented by the window pair having highest mutual writhe found among all pairs with positive (negative) mutual writhe values; this is the search tool corresponding to the unrestricted search. It searches for and ranks the structures containing at least one particular positive (negative) mutual writhe case above the threshold, independently of structure length. In the other version of rar0 (B) we use an average score of all pairs having an absolute mutual writhe above the set threshold, the average being over all pairs; this allows to search for structures having a high content of mutual writhe per squared structure length.

The two other types of scans (rar1 and rar2) use a more advanced approach based on translating the GIs into “words” or “fragment types”, by binning the GI values for each sub-chain and each sub-chain pair. Then, roughly, by word-matching with a back ground (dictionary), a query is assessed by its word/fragment-distribution: the more this differs from that found in the back ground, the higher the score. The back ground is here created by a rar1/2 run of a set of PDB-files [10] against itself, i.e. by pre-computing the GI-arrays.

More details on each scan method can be found in the Supporting Material.

### Data and implementation

We use the top100 and top8000 sets available on the Kinemage homepage [4], consisting of respectively 100 and about 8000 files of high resolution protein structures in PDB format [10].

All results are made by compiling and running the C code on a unix server, except for the computational performance (Supporting Material) which was made with a Windows-compiled version of the code run on a common laptop (Intel Core i7-4510, 2.00 GHz/2.60GHz, 8GB RAM, hard disc of SSD type; OS = win, Microsoft Windows 10).

The C source code along with implementation notes can be found in the Github repository, which also contains outlines of the code for the key functions for GISA’s tools and examples on how to run the code. The Python code and Pymol scripts for plotting and selecting examples for the restricted and unrestricted searches in this paper are also placed in the repository.

## Results

### Restricted and unrestricted searches

For the restricted search we follow [2] and let a closed loop mean a sub-chain consisting of no less than 6 and no more than 30 line-segments and such that the α-Carbon atoms at the sub-chain terminals are at most 7 Ångström apart.

In our (restricted) search for links we store the mutual writhe value of all pairs of such closed loops, while when searching for pokes keeping all candidates would result in a rather large output. To limit this, for each closed loop within a chain we keep only the two sub-chains (potential pokes) of highest and lowest writhe value, respectively. For the unrestricted search we consider sub-chains of length 30 (results for length 15 are given in Supporting Material). Here too, the output would become very large if we store the writhe for every sub-chain pair. Also, with every high writhe value pair, there will be several nearby sub-chains having almost the same high writhe.

To tackle this, we pick out for each structure the two cases (pairs of sub-chains) of lowest resp. highest writhe. Then, in the distribution of these extreme values, we examine the top-scoring cases. For both types of search we consider only pairs of non-overlapping sub-chains. More examples can be found in Supporting Material.

In the restricted search we see from Figure 1 (left) that the writhe values (almost) fit within the expected interval [-1, 1], and are distributed with rather heavy tails (for top100, see Supporting Material). In the unrestricted search (Figure 1, right) the range of writhe values becomes larger, but values rarely exceed *±*1.5 (less with a preset sub-chain length of 15, see Supporting Material). Notably, in the top100 set, the 1-links found in the restricted search were re-found in the unrestricted search; the two conspicuous cases surfacing from the restricted search through the top100 set, were both found among the top 10 absolute writhe value cases surfacing in the unrestricted search. However, the restricted search missed a 1-link in the 1dif protein’s B chain, while catching the similar 1-link in the A chain. The reason is that one of the loops in the B chain is not recognized as (almost) closed. While this can depend on the definition of an almost closed loop, the unrestricted search does not have this vulnerability and catches the 1-link in the B chain too. The additional cases of high writhe values (unrestricted search) are in general of more elaborate geometry; an example of particularly high negative writhe in the top100 set is shown in Figure 2 (right).

The same happens for the top8000 set where the 1-link cases are retained, albeit sometimes with lower rank (Supporting Material). In particular on the negative writhe side, the more complex geometries steal the top ranks and push the 1-links down the list (Supporting Material). As we shall see below, applying GISA’s basic scan tool to the top8000 set, automizing the unrestricted search, a series of 14 top-negative writhe cases were found, all highly similar to the “pseudo-knot” in Figure 2 (right), while cases appearing to be true knots surfaced as highest-positive writhe cases (Figure 3).

**Figure 3:**
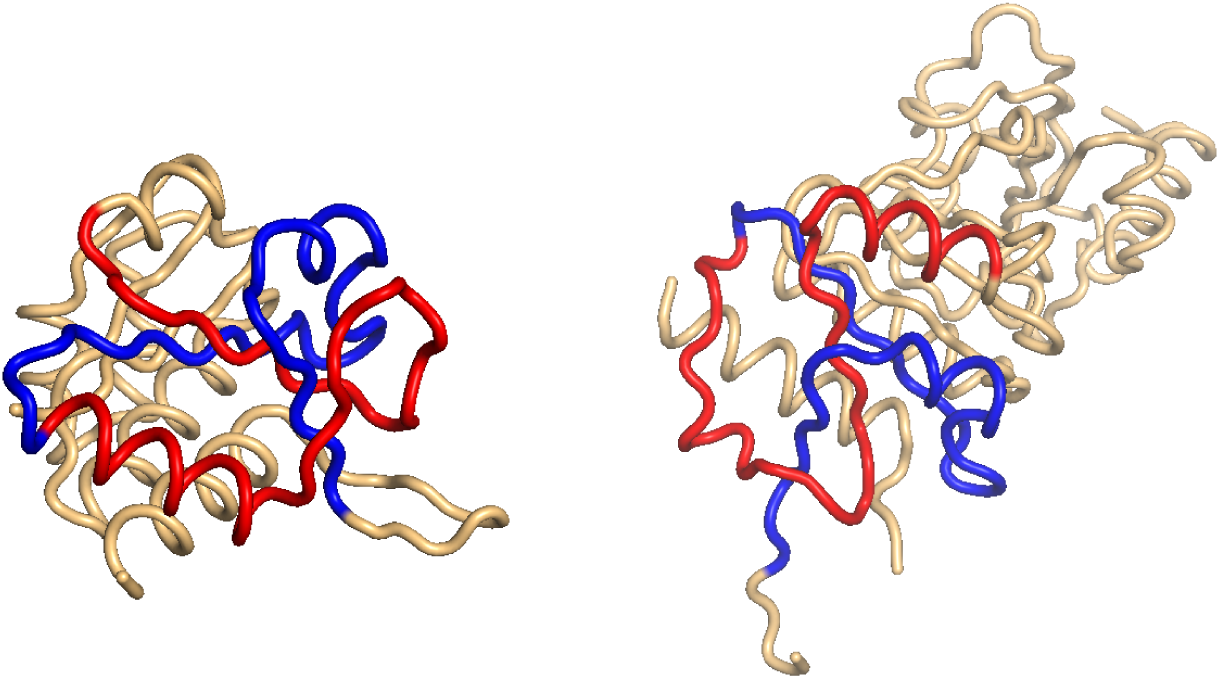
Two cases of particularly high mutual writhe value between the red and the blue sub-chains. These two knots in 3onp (left) and 2i6d (right) were found using the unrestricted search method. The plots were made with Pymol [13].

As for the pokes one may notice that the distributions of writhe values (see Supporting Material) are not heavy tailed as those for the 1-links; indeed “pokedness” is not a binary property as is linking (cf. the induction analogy above; see also Supporting Material for a short discussion). Also importantly, less pronounced writhe values reveal uninteresting examples, e.g. two loops well separated; a loop not pierced by the sub-chain (see Supporting Material).

### GISA scans

We here consider only the basic scan tool (rar0); validation of the two other tools can be found in Supporting Material. Below we show results from applying rar0 to scans of the top100 set against top8000 as back ground, and of top8000 against itself. The probability stated in the tables refer to the probability of having an absolute mutual writhe (or score) of the stated level or above among all structures in the top8000 set. For these results a sub-chain (window) length of 30 and a step size of 2 were used. The tables contain excerpts of longer lists found in the Supporting Material (Tables 7-9).

The structures found in the unrestricted search again emerge (as they should, Table 1, 2; Supporting Material Table 5, 9). We also see the clear sign of the skewness of the mutual writhe distribution and that the top100 set contains structures of rarer positive than negative writhe. In the ranking from scanning the top8000 set against itself (Table 3, 4; Supporting Material Table 7, 8) this skewness is also clear. Furthermore, in these ranks (Table 3, 4) the top 10 positive writhe cases are recorded as true knots in the KnotProt data base [5] (cf. Figure 3). At the top of the negative writhe list (rank 2-16) are 15 structures of which all except one (2v25) share a very similar structural motif of little secondary structure content (visual inspection; similar to that found in 2cpl, cf. Figure 2). Probably as should be expected, these cases are not captured in the KnotProt server [5]. Since the subchains are very similarly situated in these 14 cases, we ran a multiple sequence alignment using Clustal Omega [15] so as to grasp the sequence similarity and amount of conservation. In the resulting multiple alignment, 20 residues of the about 60 were perfectly conserved, and an additional 10 showed high similarity, an overall similarity of about 50 pct. This though does not appear as high regarding the high spatial similarity even of low secondary structural content.

**Table 1:** Top 3 structures in rar0 ranking of the top100 set vs top8000 as back ground, based on the highest positive mutual writhe pair per structure. Pair refers to the indices of the segments in the chain bordering the two sub-chains.

| Structure/chain | Pair | Mutual writhe | Probability |
| --- | --- | --- | --- |
| dif1/B | (12,42); (64,94) | 1.25 | $2.210^{-4}$ |
| dif1/A | (12,42); (64,94) | 1.25 | $2.510^{-4}$ |
| 1kap/P | (50,80);(102,132) | 1.07 | $8.310^{-4}$ |

**Table 2:** Top 3 structures in rar0 ranking of the top100 set vs top8000 as back ground, based on the highest negative mutual writhe pair per structure. Pair refers to the indices of the segments in the chain bordering the two sub-chains.

| Structure/chain | Pair | Mutual writhe | Probability |
| --- | --- | --- | --- |
| 2cpl | (70,100);(102,132) | -1.75 | $1.010^{-3}$ |
| 1nif | (230,260);(260,290) | -1.18 | $1.810^{-2}$ |
| 1php | (240,270);(270,300) | -1.04 | $4.210^{-2}$ |

**Table 3:** Top 5 structures in rar0 ranking of the top8000 set vs top8000 as back ground, based on the highest positive mutual writhe pair per structure. Pair refers to the indices of the segments in the chain bordering the two sub-chains

| Structure/chain | Pair | Mutual writhe | Rank |
| --- | --- | --- | --- |
| 3onp/A | (60,90);(106,136) | 1.50 | 1 |
| 2i6d/A | (164,194);(196,226) | 1.47 | 2 |
| 1ual/A | (74,104);(104,134) | 1.46 | 3 |
| 1ns5/B | (62,92);(92,122) | 1.45 | 4 |
| 3o7b/A | (124,154);(162,192) | 1.44 | 5 |

**Table 4:** Excerpt of 5 cases in the top15 structures in rar0 ranking of the top8000 set vs top8000 as back ground, based on the highest negative mutual writhe pair per structure. Pair refers to the indices of the segments in the chain bordering the two sub-chains.

| Structure/chain | Pair | Mutual writhe | Rank |
| --- | --- | --- | --- |
| 3hms/A | (0,30);(56,86) | -1.84 | 1 |
| 2r99/A | (70,100);(102,132) | -1.76 | 2 |
| 2wfj/A | (78,108);(110,140) | -1.76 | 4 |
| 1ixo7/B | (72,102);(104,134) | -1.72 | 10 |
| 3ich/A | (76,106);(108,138) | -1.68 | 15 |

While of relevance in itself, this case also points to using the Gauss Integrals for structural alignment; to find such structural similarities by means of superimposing the structures, if at all a viable approach, would be computationally unsurmountable (with about 100 windows in an average structure and 8000 structures it would take ∼ 10^15^ superpositions). We also noticed other cases where the mere value of the (high) writhe seems roughly to determine the spatial configuration: in the unrestricted search through top8000 at sub-chain length 15 all the top 10 ranking cases (either sign) were of the same “pseudo 1-link” nature (Supporting Material).

## Discussion

While there are obvious advantages of using a method not restricted to a sought-after geometry, such as the ability to find new configurations in real proteins and identifying non-protein like ones in models, there is a price to the generality: the unavoidable loss of specificity. In our method this shows up in the loss-of-rank for the 1-links in the unrestricted search through the top8000 set, though this essentially only hits the negative writhe cases. However, with a restricted method there is, in addition to its specificity, a likewise unavoidable weakness given by the fact that a definition of the sought-after shapes must be implemented; we saw how one of two highly similar 1-links in the top100 set was missed in the restricted search because a sub-chain was not qualified as a loop.

Regarding the rareness of the shapes (within the considered data sets), the distribution of the writhe for potential links in the top8000 set (Figure 1, left) is clearly heavy tailed. Among these around 1.4 million potential links, only about 1 out of 100000 has a writhe below −0.906 or above 0.945, respectively, i.e. there are about 28 such cases in these *∼* 8000 structures. Among the potential pokes in these structures, about 1 out of 10000 has a writhe below −0.880 or above 0.900, amounting to about 48 of such cases. It should be noticed that these numbers relate only to rareness within the top8000 set and not to whether this is a representative set of all proteins or not. However, for the links the level compares well to that reported in [2], where 37 links (“co-pokes”) were found in about 10000 real proteins. Regarding the pokes, the numbers cannot be compared; first poking may be more or less – it is not a binary topological property but rather geometric, and therefore continuously graded; second the method in [2] involves a filter which sifts out most but not all pokes in real proteins. Regarding the results in [8] these all pertain to configurations involving loops closed by covalent bonds (e.g. a cysteine bridge), and so are incommensurable to the results reported here. Identifying such configurations rather suggests a separate application of GISA in conjunction with the amino acid sequence.

Finally, the computational complexity of GISA’s base algorithm for computing the GIs of order less than three is *O*(*L*^2^), while *O*(*L*^3^) in order three (*L* being the length of the chain). This implies that getting the additional local GI values does not result in a severe time consumption. In test runs the performance of GISA compared well with that of the algorithm in [14]. This is remarkable since when run in order three (or order two “full” mode) the output of GISA even includes the GIs of order two on all connected sub-chains. At these settings the output also includes order three (resp. order two) invariants of relative nature for the sub-chains; these invariants contain information about the sub-chain and how it sits in the surrounding chain. When run in order one (and beyond) GISA produces the order one GIs on all connected sub-chains, where in our implementation we find a run time of ≈ 2 10^*-*7^s *L*^2^. The additional time spend on computations done for the proof-of-concept searches amounts to a small overhead of less than 5% (see section Computational performance in Supporting Material for these matters). The free availability of our code should make it feasible to make comparisons of timely performance of other methods to that of GISA.

## Conclusions

We have shown that with the help of GISA it is possible to find cases of rare geometries in proteins, such as those studied in [2] and knots as identified with KnotProt [9]. GISA’s command line tool scans formalize this, and more generally scores all the involved structures. The basic rar0 scan corresponds to the approach in the unrestricted search.

The method allows unprejudiced searching, in which other more elaborate shapes are found, while still catching the interesting cases found in the restricted search. Unavoidably, some specificity is lost. As such, the method shows the advantage of quantitative topological fold descriptors. Here the focus has been on applying the lowest order GI (the writhe) and a local search; GISA covers higher order GIs and supports the full range from local to global analysis, which we intend to exploit in upcoming work. In one direction, the two more advanced scan methods can be seen as foundation for making structural alignments in the space of Gauss Integrals.

## Availability of data and materials

The data that support the findings of this study are available at the Kinemage homepage, [4]. The C-code used for generating the results of this study is available under GNU General Public License v3 at GitHub (https://github.com), repository ceegeeCode/GISA (doi 10.5281/zenodo.1038274). Auxiliary Pythoncode for generating the plots is available at the same site.

## Authors’ contributions

The work was initiated in cooperation between the authors. CG created the algorithm and implemented GISA, and suggested its general local applicability. PR suggested the concrete application to identification of pokes and links. CG ran the code and wrote the first version of the manuscript; all three authors were involved in writing and revising the final version.

## Supporting Material

File “Supplementary material” in pdf format.

## References

[1] D. Zhi and M. Shatsky and Steven E. Brenner (2010). Alignment-free local structural search by writhe decomposition. Bioinformatics, 26, 1176–1184.

[2] F. Khatib and C.A. Roll and K. Karplus (2009). Pokefind: a novel topological filter for use with protein structure prediction. Bioinformatics, 25, 281–288.

[3] Jarmolinska A.I. et al. (2018). GapRepairer: a server to model a structural gap and validate it using topological analysis. Bioinformatics, 34, 3300–3307.

[4] Kinemage (2016). kinemage.biochem.duke.edu.

[5] KnotProt (2019). https://knotprot.cent.uw.edu.pl/.

[6] Marks D. et al. (2011). Protein 3D Structure Computed from Evolutionary Sequence Variation. PLoS ONE, 6(12).

[7] P. Røgen and B. Fain (2003). Automatic classification of protein structure by using gauss integrals. PNAS, 100, 119–124.

[8] P. Dabrowski-Tumanski and J.I. Sulkowska (2017). Topological knots and links in proteins. PNAS, 114, 3415–3420.

[9] P. Dabrowski-Tumanski et al. (2019). KnotProt 2.0: a database of proteins with knots and other entangled structures. NAR, 47, 367–375.

[10] PDB (2016). http://www.rcsb.org.

[11] P.L. Chang and A.T. Rinne and T.G. Dewey (2006). Structure alignment based on coding of local geometric measures. BMC Bioinformatics, 7, 346–356.

[12] P. Røgen and H. Bohr (2003). A new family of global protein shape descriptors. Math. Biosciences, 182, 167–181.

[13] Pymol (2016). The PyMOL Molecular Graphics System, Version 1.8.2.0.

[14] Røgen, P. (2005). Evaluating protein structure descriptors and tuning Gauss integral based descriptors. J. Phys.: Condens. Matter, 17, S1523–S1538.

[15] Sievers, F. et al. (2011). Fast, scalable generation of high-quality protein multiple sequence alignments using Clustal Omega. Molecular Systems Biology, 7:539.

