## Supporting Material for "GISA: Using Gauss Integrals to identify rare conformations in protein structures"

#### 1 Overview

We first provide more details on our new algorithm, GISA, for the computation of the Gauss Integral invariants [6]. We start out with the very definition of the invariants and provide a "template equation" for the recursion. From this a quite remarkable key feature of the algorithm follows easily: it is possible to compute the invariants with only little extra time consumption[1] in a recursion, which computes not only the invariants of order 1, 2 and 3 on the full structure, but gives simultaneously the values of invariants on all its sub-chains[2]. It is this "richness" that allows the present application.

Below we explain the recursion method for the lowest order case (viz. the writhe) and treat the higher order invariants by only giving the details in two cases. Thereafter we give a more comprehensive set of examples of the searches for "links and pokes" than the few found in the main paper; we show results from both the top100 set and the top8000 set [2] including 3d-plots of conspicuous cases. We then turn to the rarity scans, but only briefly, to describe a few tests and to show examples from two runs. Finally, the computationally performance of GISA is treated by some concrete examples.

[1] As mentioned in the section on computational complexity of the main paper, in test runs GISA ran at a pace comparable to the algorithm of [7]; the ratio in time consumption was between 1 and 2.

[2] To get the order 2 GIs on all connected sub-chains, GISA must be run in order 3 or order 2 "full" mode. In order 2 and 3 additional sub-chain values of invariants of a relative nature are had, but this is of no further relevance here. For more about the output of GISA see the section Computational Performance

#### 2 Method

The 29 measures (or: invariants) introduced by [6], can be computed from the 3d-structure of a fold by working out values of so called Gauss integrals. The computation of these integrals is made possible by the powerful Gauss-Bonnet theorem [6]; representing the $\alpha$-Carbon trace as a sequence of line segments (a polygonal curve, a piece-wise linear curve), the measures can be formulated in terms of sums of writhe-contributions for pairs of the segments [6]. As for defining formulas and interpretation of the invariants see [6]. In this note we use a similar notation.

##### 2.1 The algorithm

To define the invariants consider a particular fold represented as a polygonal curve in 3d-space. This consists in an ordered set of line segments, $\{s_i\}$, $i = 1, \dots, L-1$, connecting neighbouring C-$\alpha$'s, where $L$ is the number of $\alpha$-Carbon atoms (C-$\alpha$'s) in the protein's back bone. The first, and fundamental, invariant is the *writhe* of the curve [6]

$$I_{12}(1, L-1) \quad = \sum_{1 \leq i < j \leq L-1} w(i,j)$$

where each $w(i,j)$ is the writhe contribution for the pair of segments $(s_i, s_j)$ [3]. These contributions can be computed by means of the Gauss-Bonnet theorem, cf. [6]. For our purpose it suffices to know that the $w$-terms (the $w(i,j)$'s) can be computed using the coordinates of the end points of the segments. Some additional notation is handy: we refer to the set of index pairs $\{(i,j) | i < j; i,j = m, \dots n\}$ as the *simplex below* $(m,n)$ or just the *simplex* in case $(m,n) = (1, L-1)$; the elements in a simplex we refer to as *vertices*. The reason why the summation above can be taken to be only over the simplex is that $w(i,i) = 0$ and $w(i,j) = w(j,i)$ for all relevant values of $i$ and $j$ (clearly, we also have $w(i,i+1) = 0$: two consecutive line segments always sit in a plane).

We will consider the writhe of sub-chains too; for the sub-chain consisting of the segments $m$ to $n$ we simply let

$$I_{12}(m,n) \quad = \sum_{m \leq i < j \leq n} w(i,j)$$

[3] The writhe referred to in the main text is defined as $\frac{1}{4\pi} I_{12}$ as is customary; we omit the normalization here and follow the notation of [6]

An absolute version of the writhe, denoted $I_{|12|}$, is defined by summing up the absolute value of the writhe contributions, $|w(i,j)|$. While the writhe computes a signed average crossing number, $I_{|12|}$ computes its unsigned companion, the average crossing number (see [6] for more on the interpretation).

The recursion equation for the writhe, which is at the same time the template for all the higher order cases, is arrived at by noting that $I_{12}(m,n)$ satisfies a very simple decomposition illustrated here (Fig.S1) for a sub-chain consisting of the segments from $m = 50$ to $n = 90$:

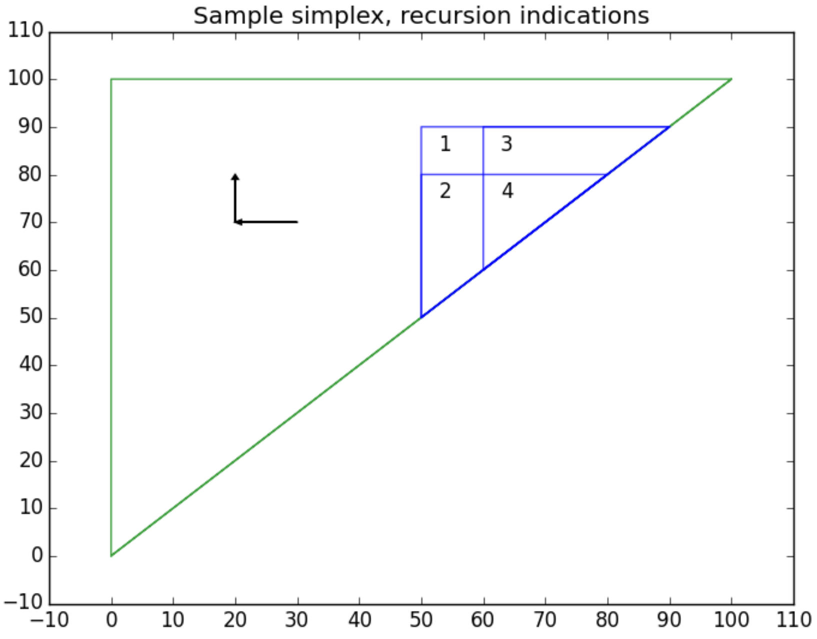


Fig.S1. Example of the simplex (all vertices within the green border) for a chain of 100 segments.

In this example we can write

$$\begin{aligned} I_{12}(m=50, n=90) &= \text{Area1} + \text{Area2} + \text{Area3} + \text{Area4} \\ &= \text{Area1} + (\text{Area2} + \text{Area4}) + (\text{Area3} + \text{Area4}) \\ &\quad -\text{Area4} \end{aligned}$$

The point with this is that the last three terms are writhe values of the sub-chains corresponding to the areas (here the sub-chains 50-80, 60-90 and 60-80 respectively). In particular, if we just consider the sub-chains given by removing the C-alpha at the beginning of the $m-n$ sub-chain or at its end, we have

$$I_{12}(m,n) = \\ w(m,n)+I_{12}(m+1,n)+I_{12}(m,n-1)-I_{12}(m+1,n-1) \quad (1)$$

As this formula suggests, if we implement the computation of $I_{12}(1,L-1)$ in a 2d for-loop over the simplex, moving backward in the values of $m$ (from $L-1$ and down to 1) and, for each such $m$, moving forward in the value of $n$, from $m$ to $L-1$, we can compute $I_{12}(m,n)$ from $w(m,n)$ and already known values of $I_{12}$ (as indicated by the arrows). Indeed, the right-hand side can be computed if we know a) "the corner value" $w(m,n)$ and b) the values of $I_{12}$ at vertices $(i,j)$ *prior* to $(m,n)$ in the sense that $(i,j)$ has been passed in this 2d-loop when it arrives at $(m,n)$ (that is, if $i>m$ or if $i=m$ and $j<n$). We may sum this up by saying that the computation of $I_{12}$ at any vertex $(m,n)$ can be done in a recursion *by prior vertices and measures* — regarding, with sense, $w$ as a measure prior to $I_{12}$. We notice that it is clear that the same recursion formula and statements hold for $I_{|12|}$, though we shall not make use of this here.

Importantly, we can here see how the "richness" property alluded to above emerges. We ultimately want to know the value of $I_{12}$ on the whole chain, i.e. the number $I_{12}(1,L-1)$. However, implementing the computation of $I_{12}$ as we have just outlined will provide us with the value of $I_{12}$ on all sub-chains of the given chain. Thus, the recursion fills out the simplex with values of the invariant. This richness we shall exploit here to compute swiftly the *mutual writhe* of any two given sub-chains of a larger chain; this is the sum of writhe contribution stemming from segment pairs having one segment from each chain (i.e. the "mixed" contributions). For the searches that we are aiming for, the mutual writhe is exactly what we need, since it amounts to the linking number of the two sub-chains.[4] Let us here understand this notion in the setting of Figure S1: there the mutual writhe of the sub-chains (50,60) and (80,90) is the sum of the $w$-terms in Area1 (including the boundary). In the equation right below the figure and taking the areas appropriately without the boundary that now goes into Area1, as we clearly may, the left-hand side is the writhe of a sub-chain and on the right-hand side the last three summands are also writhe values of sub-chains. What remains is Area1, which can therefore be obtained from writhe values of sub-chains (as a simple signed sum).[5]

[4] While an ideal loop has zero writhe, this will conveniently also disregard any writhe each sub-chain may have with itself.

[5] In [6] and [7] this was used to half the computational complexity of the higher order

##### 2.2 Recursion for higher order invariants

While the recursion method does not improve on the performance of the computation of the writhe, this is not the case for the higher order invariants. However, since not the primary focus of our search method, we dwell on them only briefly and give details in two cases. The higher order measures are sums of products of $w$-terms, with the order referring to the number of factors ($w$-terms) in the products. For instance [8]

$$I_{132645}(1, L-1) \quad := \sum_{1 \leq a < b < c < d < e < f \leq L-1} w(a,c)w(b,f)w(d,e)$$

defines an order 3 invariant (while the summation order is 6 it can be brought down to 3; this follows from the present algorithm and is well-known [8]). Following [4] we consider these invariants up to and including third order and, as it turns out, all these fit into recursion formulas of the same shape as that for the writhe. Suggestively, for invariant $I$ we can write

$$\begin{aligned} I(m,n) \quad &= \quad \text{fct (lower order invariants at (m,n) or prior vertices)} + \\ &\quad\;\; I(m+1,n) + I(m,n-1) - I(m+1,n-1) \end{aligned}$$

where $w$ will be understood as a lower order invariant for any given $I$. We refrain from carrying out the detailed derivation, which for each invariant reveals what the function term "fct" is. For the sake of inspiration though let us consider the simple order 2 invariant, $I_{1423}$. By definition [8]

$$I_{1423}(m,n) \quad = \sum_{m \leq i < a < b < j \leq n} w(i,j)w(a,b)$$

Then clearly

$$I_{1423}(m,n) \quad = \sum_{m \leq i < j \leq n} w(i,j) I_{12}(i+1, j-1)$$

So writing

$$w_{1423}(i,j) \quad = \quad w(i,j) I_{12}(i+1, j-1),$$

we see that $I_{1423}(m,n)$ is a sum of the $w_{1423}$-terms over the simplex below $(m,n)$ (i.e. the simplex which has $(m,n)$ as top-left corner) and since these

Gauss Integrals.

$w_{1423}$'s only depend on these vertices the case is as with $I_{12}$. It follows that we have the recursion equation

$$\begin{aligned} I_{1423}(m,n) &= \sum_{m\leq i<j\leq n} w_{1423}(i,j) \\ &= w(m,n)I_{12}(m+1,n-1) \\ &\quad +I_{1423}(m+1,n)+I_{1423}(m,n-1)-I_{1423}(m+1,n-1) \end{aligned}$$

We see that indeed the first summand, $w(m,n)I_{12}(m{+}1,n{-}1)$, is a "function of lower order invariants at $(m,n)$ or prior vertices" and conclude that $I_{1423}$ can be handled in the same 2d for-loop as $I_{12}$.
The derivations of the recursion formulas (i.e. to arrive at transparent "fct"-terms) do not go quite as smoothly for all invariants, but at any rate they are not very difficult to arrive at. In a few cases new measures playing the role as fct-term pop up ("relative" invariants) and in a couple of cases several such measures emerge, and a canceling out among these leads to simpler expressions. The recursion formulas show that we can compute the majority of the measures of order 1, 2 and 3 in a 2d for-loop. Unsurprisingly, a handful of the order 3 measures though seem to be out of reach: they call for a 3d-loop – or, put differently: they are polynomials in the $w$-terms of an order strictly larger than 2.
To end this section let us pause to show how a relative measure shows up in the recursion formula for $I_{1234}$. By definition [8] we have

$$I_{1234}(m,n) = \sum_{m\leq i<j<a<b\leq n} w(i,j)w(a,b)$$

So clearly

$$\begin{aligned} I_{1234}(1,L) &= \sum_{1\leq i<j\leq L} w(i,j) \sum_{j<a<b\leq L} w(a,b) \\ &= \sum_{1\leq i<j\leq L} w(i,j)I_{12}(j+1,L) \end{aligned}$$

The right-hand side here has the same shape as in the definition of $I_{12}$. So if we write

$$w_{1234}(a,b;c) := w(a,b)I_{12}(b+1,c)$$

and define a relative version of $I_{1234}$ by

$$I_{1234}(i,j;l) \quad := \sum_{i \leq a < b \leq j} w_{1234}(a,b;l)$$

we have $I_{1234}(1,L) = I_{1234}(1,L;L)$ along with the recursion formula

$$I_{1234}(i,j;L) =$$
$$w_{1234}(i,j;L) + \quad I_{1234}(i+1,j;L) + I_{1234}(i,j-1;L) - I_{1234}(i+1,j-1;L)$$

##### 2.3 GISA's scan methods: scoring and more

This section is dedicated to the details of the rarity scan/detection functions of GISA. As explained in the main paper, rar0/1/2 are functions for ranking one or more structures (queries) by comparison to a back ground "data base" (based on a set of PDB files). In particular, this allows assessing to what extent the queries stand out as "rare" on that back ground. The code allows running one or more sets of queries against the data base without reloading the back ground (which includes memory allocation for loading the GI's for the complete back ground PDB-set; for rar1/2 the "dictionarying", i.e. translation of the GI-arrays into words by binning the values; and sorting the values/words).

The following sections explain how each of the scans work, including their scoring.

###### 2.3.1 Flavour: rar0

In rar0 rank is decided by means of mutual writhe (possibly absolute value thereof, but signed by default). The method is as follows (see also the examples runs in the Github repository):

1. Create the back ground "data base" of the wanted Gauss numbers for pairs of windows (only needed if using version B below)

2. Run the rarity scan with rar0: there are two methods of scoring available:

   A Score by the maximal mutual writhe: for a given query (q) we pick out the highest absolute mutual writhe among all the positive (negative) writhe values in the structure, or, if running in "unsigned mode" we just keep the pair for which the absolute value of the mutual writhe is highest (so that in this unsigned mode there is one value, max-abs-mutual-writhe, for each q, while in the signed case we consider the highest positive and the highest negative). The obtained value is then held up against the distribution of similarly obtained max mutual writhe values across the structures in the (background) data base (signed: two distributions, one for positive writhes and one for negative; unsigned: a single distribution of max-abs-mutual-writhe values). This provides directly a "p-value" (and a score = $-log(p)$-value), viz. the frequency of max (abs) mutual values in the data base more extreme than the obtained value (in particular step 1 above is not needed in this mode).

B Score by absolute mutual writhe above a set threshold: for a given query (q) we run through all pairs of windows in q; for each pair for which the absolute mutual writhe ($amw$) is above a set threshold, T (e.g. 5), we find the probability that a pair in the database has an absolute mutual writhe higher than this absolute mutual writhe ($amw$) for the given pair of windows (i.e. probability(abs mutual writhe > $amw$)). This is just a look-up in the background distribution of absolute mutual writhes. The final score for the query is then the sum of $-log(p)$-values in this set of pairs

$$score(q) = -\sum log(p)$$

where the sum is over all pairs in q with $amw > T$ as just explained. The final score is now the average of this score, i.e.

$$score = -\frac{\sum log(p)}{\#\text{pairs in query}}$$

where # means size of). It is possible to use the non-averaged scored as final score (average score is default).

A reason for using the average score (in B) is that the probability of having some (rare) 3d-configuration should increase with the length of the structure (here quadratically as the number of pair is quadratic as a function of the length of the query). However, using the average score has some disadvantages, e.g. a short structure with one rare window pair will get a higher score than a longer structure with exactly the same odd pair and no other particularities; on the other hand, such short structures ought to appear more rare if "weird 3d-configurations" are distributed uniformly over all pairs in the database. In addition version A allows getting the score based on the most extreme pair rather than just an average consideration (and, in addition, in the unsigned mode).

With version B, to obtain a final p-value corresponding to the obtained score, it is necessary to first run rar0 with query set = data base and with absMutValScorePValues_b = 0 (i.e. to carry out step 1 above). This generates the background distribution of scores (and of amw's); if we believe that the data base consists of a representative set of structures (for a given purpose), we can with reason score any query set against this background. So when this run for a back ground is done, we run rar0 for the desired query set now with absMutValScorePValues_b = 1. In version A, p-values and corresponding scores are had directly (so, as mentioned, step 1 is not needed).

The scoring method A scans for structures having one (or more) exceptional mutual writhe pairs and allows distinguishing between the positive writhe pairs and the negative writhe pairs; the B version scans for structures having possibly several pairs of high absolute mutual writhe (above the threshold) but maybe none of an exceptional level. With B a high threshold on the mutual writhe should be used, e.g. 10.

###### 2.3.2 Flavour: rar1

While rar0 only uses the mutual writhe, in the flavours rar1 and rar2 it is possible to use all invariants up to and including order two; therefore in these flavours the GI values will be arrays/tuples of the individual GIs (the length of the array is the number of GIs chosen for the scan).

In rar1 each window pair of a query is scored by first translating its GI-array into discretized versions — "words" — by binning the individual GIs, and then counting its matches in a likewise translated back ground (i.e. what can be seen simply as a look-up of the word in a back ground dictionary). Here a number of mismatches can be allowed. These can though only be "one bin off", i.e. in a mismatch only neighboring letters are allowed: if e.g. three invariants are used, and a GI-array is translated into ABC, AAC will be an allowed mismatch (of one), while ABA is not since the last A is two letters away from C.

In rar1 only pairs of sub-chains are considered, so only "mutual GIs" are used. However, in order two the mutual value as computed in GISA will in general depend on the chain between the sub-chains of the pair, which is inappropriate for the word-matching. Therefore in rar1 only the two first order GIs should be used. As in rar0, a threshold on the absolute value of the mutuals allows to focus on occurrence of ”more rare words” (and to lower computation time).

Here follows an outline of the method:

1. The data base is converted to a ”dictionary”: each data base element is a tuple/array of Gauss numbers (in number as many as the desired number of invariants for matching, and in any case limited the number of invariants of order 1 or order 2); each of these tuples is converted by binning the Gauss numbers into a tuple of integers (ie a ”word”); after sorting these words lexicographically the data base has the guise of a dictionary (though probably with many words repeated).

2. A given query is similarly translated, by the same binning, into a set of words (one for each window pair); the window pairs are now looped through and each is looked up in the data base dictionary (by setting a threshold as mentioned, only pairs having a mutual writhe (or mutual invariant) in absolute value above this threshold are considered).

In the matching a set number of mismatches can be allowed (as explained above too). The look-up of the query ”word” in the data base gives a count of the number of matches, $cntMatch$, for each pair. The score is now the average

$$Score = -\frac{\sum_{\text{pair in query}} log(\frac{cntMatch(pair)}{\#\text{data base}})}{\#\text{pairs in query}}$$

which is the same as

$$Score = -\sum_{\text{word w}} \frac{\#\text{pairs in query of word w}}{\#\text{pairs in query}} log(\frac{\#\text{db-pairs of word w}}{\#\text{data base}})$$

If we write $p_q(w) = \frac{\#\text{pairs of word w}}{\#\text{pairs in query}}$ and $p_{db}(w) = \frac{\#\text{db-pairs of word w}}{\#\text{data base}}$ we then have

$$Score = -\sum_{\text{word w}} p_q(w) log(p_{db}(w)),$$

a cross entropy that is, and also showing the resemblance to the Kullback-Leibler relative entropy

$$KL = \sum_{\text{word w}} p_q(w) log(\frac{p_q(w)}{p_{db}(w)})$$

i.e. only the "q-idiosyncratic" entropy term "$\sum_{wordsw} p_q(w) log(p_q(w))$" is disregarded.

The rar1 scoring method ranks the structures on their "distribution of words" as compared to that of the back ground, their significance increasing with the score. One can also think of this as a way of determining whether a query has an unusual set of fragment pairs as compared to the background. The threshold allows to focus on occurrence of "rarer words".

##### 2.3.3 Flavour: rar2

While rar0 and rar1 only use pairs of sub-chains, rar2 is based on single sub-chain (window) matching, but which can be added a pairs-based scoring. The single window matching works by translating the GI-arrays into words as in the pairs case (rar1). The optional pairs matching is done based on the single window matches: considering a window pair in the query structure, the matches in the data base are those among the corresponding set of pairs of matching single windows, for which the mutual GIs also match those of the query. Here a number of invariants and a number of allowed mismatches can be set both for the single and the pairs window matching; while the single window matching can be done including order two GIs, as in rar1 it is recommendable in the pairs matching to only use the order one invariants. As with rar0/1 a threshold allows to focus on occurrence of "more rare words" in the pairs matching part.

As in rar1 the scoring in rar2 is done by means of a "cross entropy". Here follows an outline of the method for the single window matching:

1. The data base is converted to a "list of words": each data base element is a tuple/array of Gauss number (in number as many as many as there are invariants of the chosen order, ie 1 or 2); each of these tuples is converted by binning the Gauss numbers into a tuple of integers (i.e. a "word"); after sorting these words lexicographically the data base has the guise of a dictionary (though with probably many words repeated). As opposed to rawRarity1 this "dictionarying" is based on the Gauss numbers for the windows and not the window pairs.

2. A given query is similarly translated into a set of words (one for each window); the windows are now looped through and each is looked up in the data base dictionary (for a pairs-based scan see more right below). In this matching, a pre-set number of mismatches can be allowed (as in the rar1 pairs-based version).This gives a count of the number of matches, cntMatch, for each window.

Again, as in rar1, the look-up gives a count of the number of matches, $cntMatch$, for each pair. The score is now the average

$$Score = -\frac{\sum_{\text{window in query}} log(\frac{cntMatch(window)}{\#\text{data base}})}{\#\text{windows in query}}$$

which, just as in rar1, but with "windows" rather than "pairs" can be rewritten as

$$Score = -\sum_{\text{word w}} \frac{\#\text{windows in query of word w}}{\#\text{windows in query}} log(\frac{\#\text{db-windows of word w}}{\#\text{data base}})$$

If we write $p_q(w) = \frac{\#\text{windows of word w}}{\#\text{windows in query}}$ and $p_{db}(w) = \frac{\#\text{db-windows of word w}}{\#\text{data base}}$ we then have

$$Score = -\sum_{\text{word w}} p_q(w) log(p_{db}(w)),$$

a cross entropy that is, resembling the Kullback-Leibler relative entropy as in rar1 above.

If a scan based on the pairs is also wanted, the pairs of windows in a query will subsequently be looped through (i.e. after all windows of the given query have been looped through and the matches are found and recorded); it is possible and desirable for speed to set a threshold so that only pairs having a mutual writhe (or mutual invariant) value above this threshold are considered). The scoring is verbatim the same as in the single window case, simply replace window by window pair (or see rar1).

The rar2 scoring method scans for structures having a "distribution of words" significantly different from that found in the background distribution. One can also think of this as a way of determining whether a query has an unusual set of fragments/fragment pairs as compared to the background. The threshold allows to focus on occurrence of "more rare fragment paris" in the pairs-based matching.

### 3 Results

In this section we supply additional results from searching the top100 set and the top8000 set [2]. For the restricted search we include results from both links- and pokes-searching; for the unrestricted search there is naturally no such distinction: the geometries/configurations are rather output of the search.

#### 3.1 Restricted search

We have explained in the main text how to search for links of (almost) closed loops by means of the mutual writhe and similarly for the pokes. The intuition is as mentioned to think of the electromagnetic induction in a wire poking through a loop of another wire in which an electric current runs. We may then think of our search as one for cases of exceptional induction. In what follows we generally suppress the word mutual.

We first consider the top100 set. For the run we used the settings stated in the main text: min and max loop lengths of 6 and 30, respectively, and a distance of 7 Ångström for defining "closed". Here (Fig.S2) is then first the distribution of the writhe values for all potential links (i.e., pairs of closed loops in each structure) and pokes:

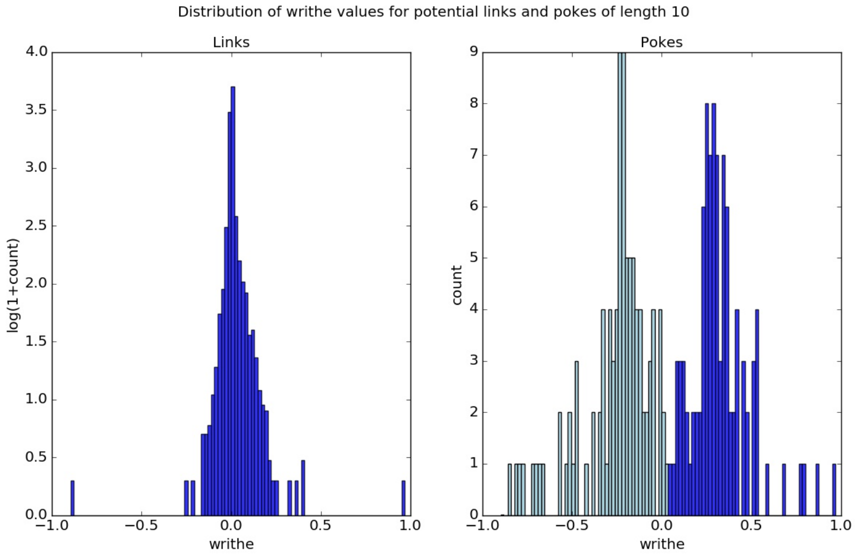


Fig.S2. Distribution of writhe values for potential links (left) and pokes (right) in the top100 set. For the pokes the distribution of the min-values (max-values) is light blue (dark blue). Please note that the counts are replaced by $log(1 + count)$ for the links. The poke-length was set to 10; other settings were as stated in the text.

As is apparent the writhe values for the links lie in a range bounded below at about -0.9 and above at about 0.95. Quite remarkably, two cases [6] stand out, their writhe in absolute value being close to 1 (the value can

[6] We disregard chains containing "holes", i.e. chain segments of a length above thresh-

diverge from 1 in either direction due primarily to other writhe contributions stemming from the shapes not being ideal, i.e. the loops are not closed; smoothing will not change the writhe except if one allows the end points of the sub-chains to move). We visualize three different orientations in each case to illustrate their three-dimensional nature. These two examples (Fig.S3a-b) and the similar 3d-plots to come are annotated as follows: the title gives the protein's name/the chain shown [7], next as two pairs of integers, the bordering residue numbers of the high-lighted sub-chains (here almost closed loops) and finally the writhe value (rounded). In the plots the red segment is "first" and the blue is "last", i.e. the red segment is the one having the lowest index range. Please note that the indication of the segments in the tables is by their indices in the chains and not by their residue numbers. Also, in the tables in the section on results from GISAs scan methods, the writhe values are as output by GISA; elsewhere here and in the main paper the values are normalized by $4\pi$ (as is customary in the definition of the linking number).

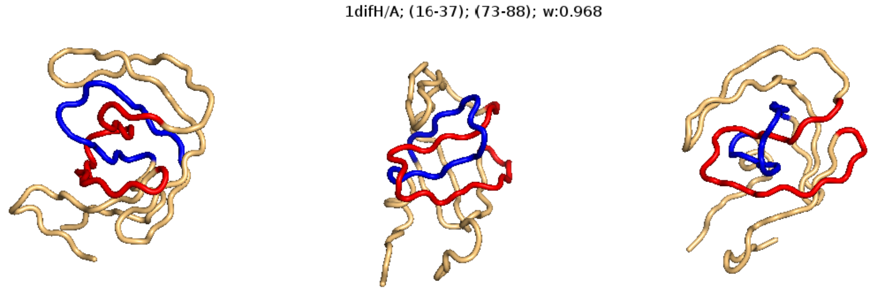


Fig.S3a. A potential link (writhe $\sim$ 0.968) in chain A of t he 1dif protein.

old, here set to 7 Ångström; the E chain of the 2er7 structure is disqualified for this reason, but contains a 1-link in parts not containing the hole.

[7] ">" means that in the PDB-file the chain id was left blank

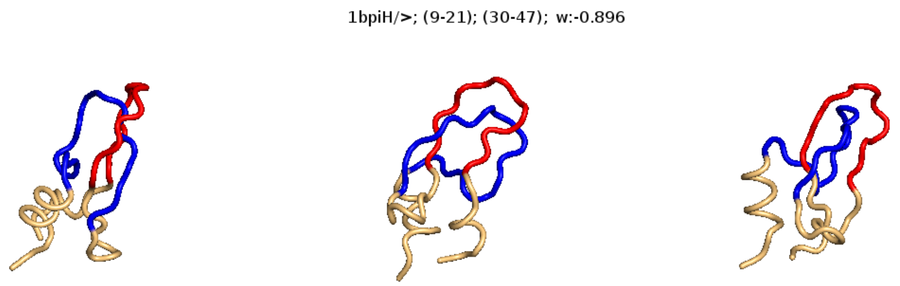


Fig.S3b. A potential link (writhe ∼ -0.896) in the rather small 1bpiH protein (length just below 60).

| Structure/chain | Pair | Mutual writhe | Type |
|---|---|---|---|
| 1dif/A | (15, 36);(72, 87) | 0.97 | link |
| 1bpi/- | (8, 20);(29, 46) | -0.90 | link |
| 1dif/B | (72, 87);(25, 35) | 0.86 | poke |
| 1phb/- | (49, 77);(281, 291) | -0.85 | poke |
| 2cpl/- | (83, 102);(119, 129) | -0.82 | poke |

Table 1: Top 2 links and the three top ranking pokes not merely part of the two links in the top100 set. Pair refers to the indices of the segments in the chain bordering the two sub-chains.

Next let us consider the search for pokes. We have used a set poke-length of 10 residues; we shall comment on this setting later. From Fig.S2. we see that the range of the writhe is bounded above at slightly below 1 and below at a little more than -1. Notably, the extreme cases are not as "lonesome" as with the links: the range is rather continuously occupied out to the highest value. One should bear in mind that for pokes we are even only keeping the highest and lowest value case for each closed loop, while for the links we are considering all candidates. Top ranking are two pokes merely part of the two links (and which are not shown in the table). Next is a poke in the B chain of 1dif; this is actually part of a 1-link similar to the one in the A chain and which the restricted search misses (but the unrestricted search captures). The poke in 2cpl is also a a part of a configuration that we show later. Here follows (Fig.S4) the poke in 1phb (there is one further example in 1phb, but the closed loop only differs by two C-alpha's from the one shown here):

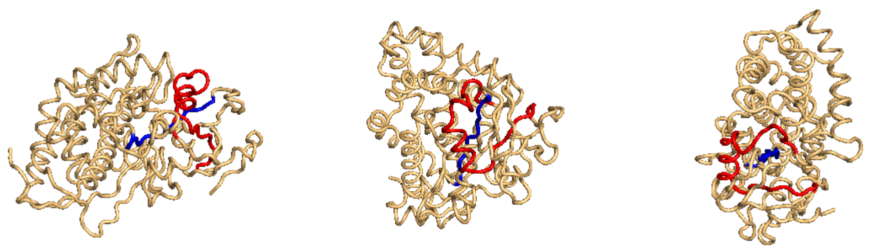


Fig.S4. A potential poke (writhe $\sim$ -0.846) in the 1phb protein.

For a poke the size of the writhe (induction) depends on the relative proportions of the poking piece and the loop, i.e. the geometry, while in an interlinking case (e.g. a 1-link) the proportions of the two are unimportant. There it is rather the threading — or: topology — which is crucial. This difference is also reflected in the distributions of writhe values for the potential links and for the potential pokes shown above (the outstanding link examples disconnected from the rest vs. the more continuously occupied range among the potential pokes). The poke examples may therefore not appear as "shining" as the links, but in each case shown here one may with reason say that the poke-segment is placed in "poking positions" in the loop. Some "sheeting" could occur, i.e. that the poke-segment is "parallel" to the loop, but this seems not to be a burden in our approach [8] (in [1] the authors apply a "sheet filter" to sift out cases of such nature).

While it is reassuring and satisfying to see that high (absolute) writhe values imply cases of truly linked loops and very qualified poke candidates, it should also be checked that at lower values the linking or poking is much weaker if at all present. Indeed, checking a few cases this appears to be the case. Here follow, for potential links, a case of medium size negative writhe (Fig.S5a), a case of writhe close to zero (Fig.S5b) and a case of medium size positive writhe (Fig.S5c); then follow two cases of potential pokes with low-to-medium writhe (in absolute value):

[8] We have noticed that if using the average crossing number, $I_{|12|}$, rather than the writhe, $I_{12}$, sheeting examples surface at the expense of "true pokes" such as those shown here.

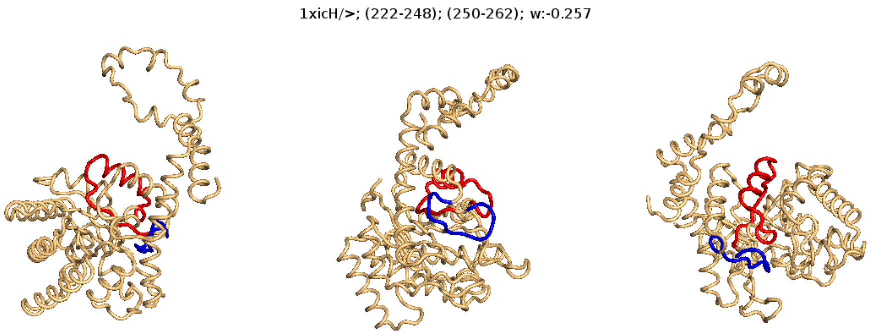


Fig.S5a. A potential link of writhe $\sim$ -0.257 in the 1xicH protein. The expectation here is that of "no-linking" as indicated by the low mutual writhe value, but probably some proximity.

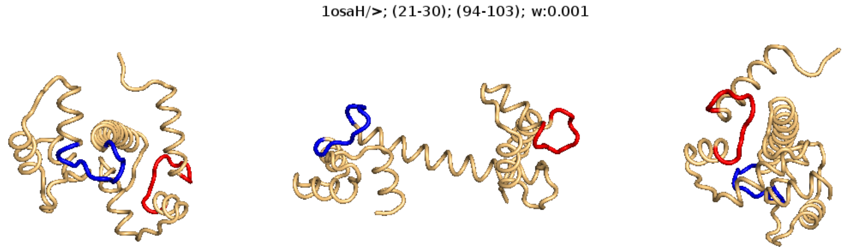


Fig.S5b. A potential link of writhe $\sim$ 0.001 in the 1osaH protein. The expectation here is that of "no-linking" as indicated by the very low mutual writhe value, as well as very little proximity.

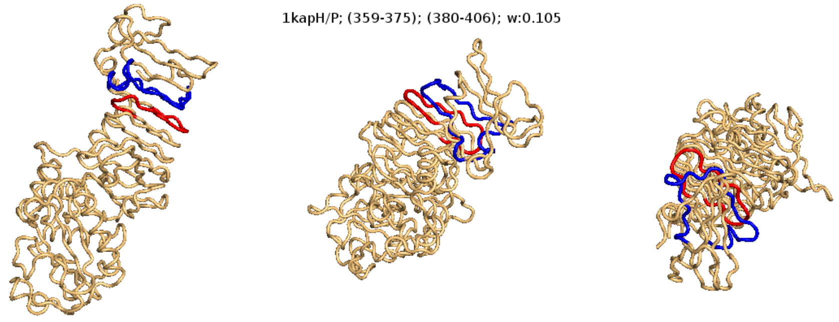

Fig.S5c. A potential link of writhe ∼ 0.105 in the 1kapH protein. The expectation here is that of ”no-linking” as indicated by the low mutual writhe value, but probably some proximity.

As expected, at a writhe close to zero the two loops are distant, while higher values arise for loops in close proximity, though not inter-linking. Next the two poke examples (Fig.S5d-e) revealing again that close proximity leads to some amount of writhe, but far from exceptional values:

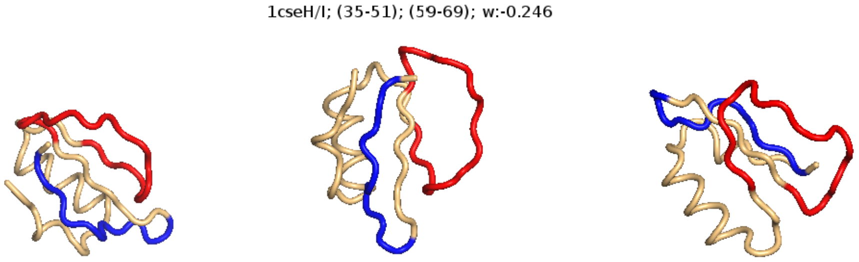


Fig.S5d. A potential poke of writhe ∼ -0.246 in the 1cseH protein. The expectation here is that of ”no-poking” as indicated by the low mutual writhe value, but probably some proximity.

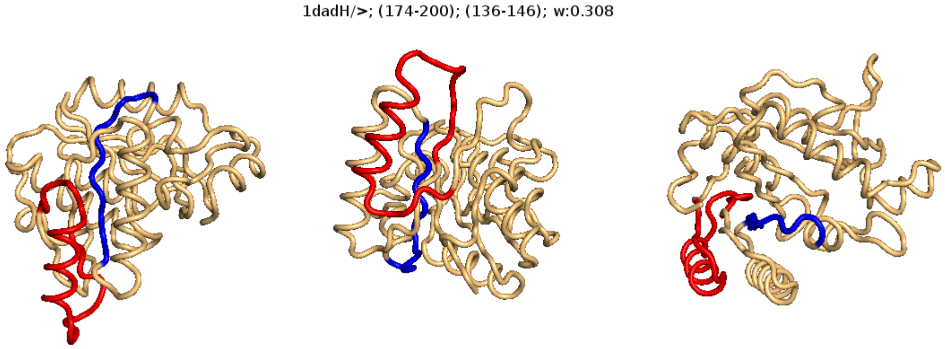


Fig.S5e. A potential poke of writhe ∼ 0.308 in the 1dadH protein. The expectation here is that of ”no-poking” as indicated by the low mutual writhe value, but probably some proximity.

So far we have only shown concrete examples from the top100 set. In the top8000 set 21 (52) cases having an absolute mutual writhe above 0.95 (0.9) were found; of these 9 (21) were of positive mutual writhe, 12 (31) negative. Here is the top5/top5 (Table 2):

| **Structure/chain** | **Pair** | **Mutual writhe** | **Type** |
|---|---|---|---|
| 1pqh/B | (9,34);(52,76) | 1.00 | link |
| 2qd6/A | (15,36);(72,86) | 0.99 | link |
| 1ual/A | (84,107);(115,139) | 0.98 | link |
| 2egv/A | (157,178);(186,212) | 0.96 | link |
| 3o7b/A | (142,161);(169,197) | 0.96 | link |
| 3hms/A | (3,31);(56,78) | -1.01 | link |
| 3dqp/A | (159,174);(188,210) | -1.00 | link |
| 3dqp/A | (159,174);(182,203) | -0.99 | link |
| 3fdr/A | (10,37);(59,82) | -0.99 | link |
| 3dqp/A | (157,171);(188,210) | -0.99 | link |

Table 2: Top 5 positive and top 5 negative writhe cases among the potential links in the top8000 set. Pair refers to the indices of the segments in the chain bordering the two sub-chains.

The remaining of the 21 cases of absolute mutual writhe above 0.95 were found in (some containing several similar/overlapping links): 2egv, 3aia, 3m3q, 2ha8 (positive writhe) and 3fdr, 3dqp (negative). The three high scoring cases look like this:

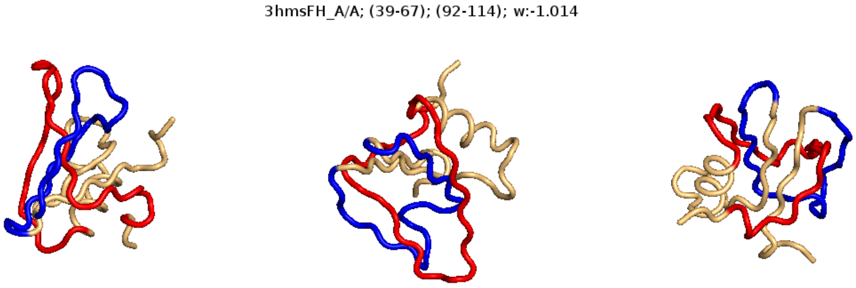


Fig.S7a. A potential link of writhe $\sim$ -1.014 in chain A of the 3hms protein.

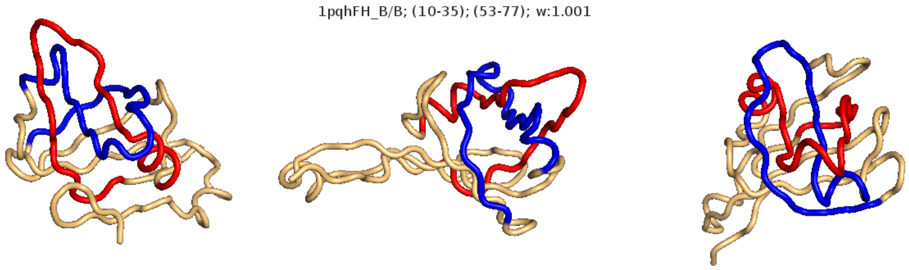


Fig.S7b. A potential link of writhe $\sim$ 1.001 in the B chain of the 1pdq protein.

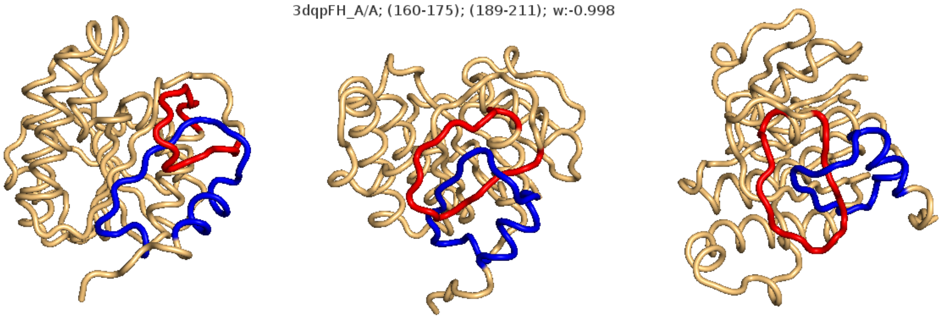


Fig.S7c. A potential link of writhe ∼ -0.998 in the B chain of the 3dpq protein.

The two other links in 3dpq are very nearby in the chain, and there are in fact more combinations of these subchains giving rise to links in this structure. The same subchains give rise to several of the high-writhe pokes in the top8000 set, which we now turn to. The highest scoring is a very clear case which we show next (Fig.S8); as we shall see later it is in fact part of a knot. Of the others only three appear merely as a part of one of the links above, viz. 2qd6, 1ual and 3dqp, while the remaining are seemingly more genuine pokes.

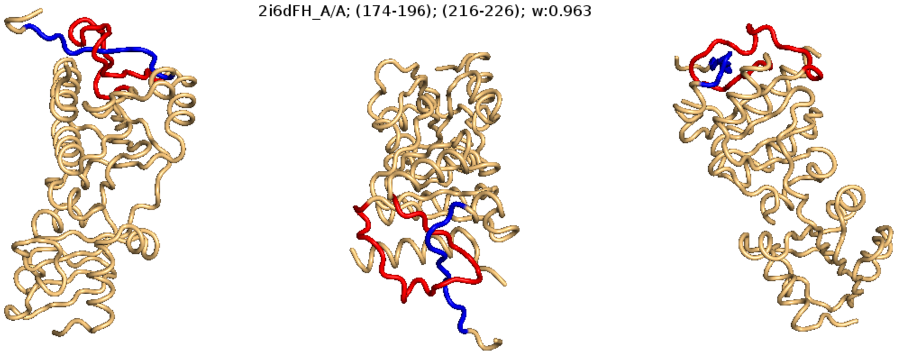


Fig.S8. A potential poke of writhe ∼ 0.963 in chain A of the 2i6d protein.

As for the set poke-length it was also tried out to use a length of 5 and

| **Structure/chain** | **Pair** | **Mutual writhe** | **Type** |
|---|---|---|---|
| 2i6d/B | (174,196);(216, 226) | 0.96 | poke |
| 2qd6/A | (15,36);(76, 86) | 0.95 | poke |
| 3m3g/A | (26,54);(72,82) | 0.95 | poke |
| 1j71/A | (210,228);(290,300) | 0.95 | poke |
| 1ual/A | (84,107);(126, 136) | 0.94 | poke |
| 3dqp/A | (159,174);(192,202) | -1.05 | poke |
| 2jh1/A | (169,199);(156,166) | -1.04 | poke |
| 3dqp/A | (157,171);(192,202) | -1.03 | poke |
| 1knt/A | (28,45);(10, 20) | -1.00 | poke |
| 3dqp/A | (153,169);(192,202) | -1.00 | poke |

Table 3: Top 5 positive and top 5 negative writhe cases among the potential pokes of length 10 in the top8000 set. Pair refers to the indices of the segments in the chain bordering the two sub-chains.

one of 7. Regarding whether any of these lengths is to prefer over the other, the distributions shed some light. To this end the top8000 results should be considered simply for its size. Here follow (Fig.S9) the writhe distributions for poke-length 5, 7 and 10:

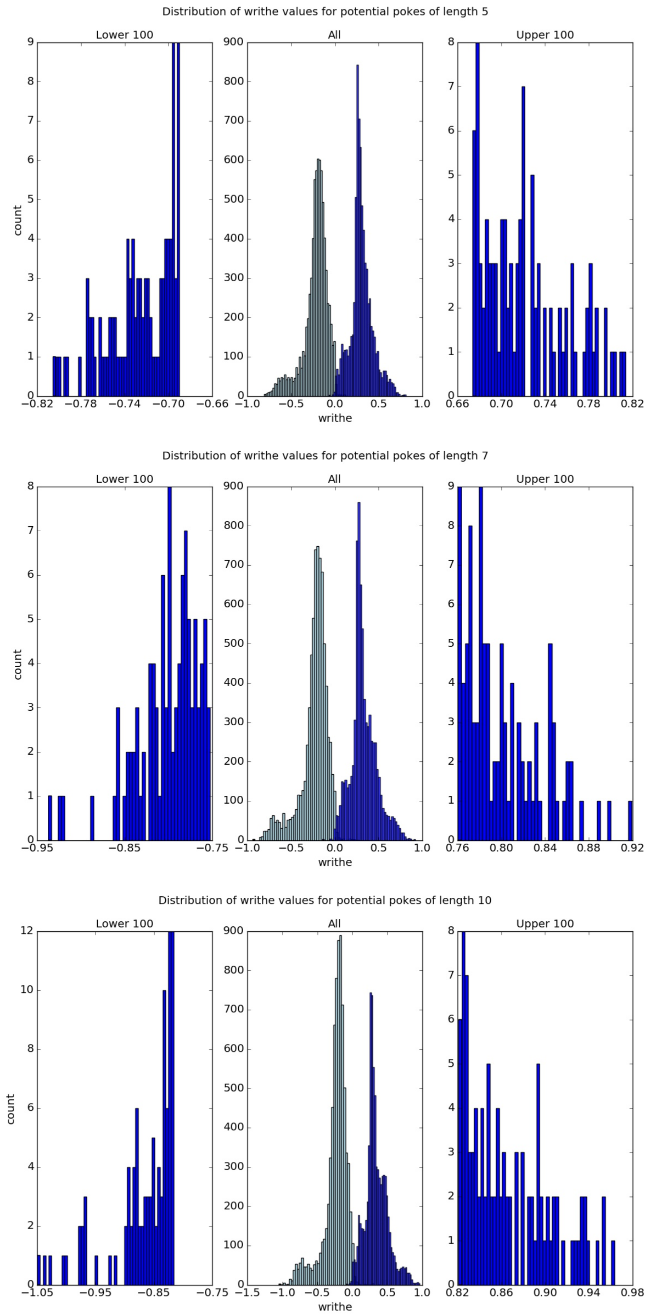


Fig.S9. Distributions of writhe for potential pokes of length 5 (top), 7 (mid) and 10 (bottom) in the top8000 set: in the middle the entire distributions of cases of lowest (light-blue) and highest (dark-blue) writhe value per chain. To the left and right a zoom-in on the tails.

With length-5 the top-scoring examples (in the left-hand tail) are rather "lonely": there is quite a gap down to the second best scoring and the potential range $[-1, 1]$ is not filled out. With length-7 the range is closing in on the theoretical one, but there is still a gap and the left-hand tail is still "thin". With length-10 the situation is improved a notch further, examples throughout the theoretical range show up, and the tails appear more connected). This could hint at that using length 10 is less fragile than the two other. On the other hand, among the top-10 examples of highest writhe value, eight are shared by the length-5 case and nine are shared in length-7 (the length-5 examples not found in length-10 are two clear pokes in the 1ra9 protein and the additional length-7 example is a likewise clear poke in 1cus). So it cannot be said that the outcome is very sensitive to the set poke-length, but still it seems most advisable to use a length of 10.

Regarding computation time, we have in the main text mentioned that the performance of the base part of the algorithm, which computes the invariants' values, is only mildly affected by adding the searches. This also goes for the unrestricted search method which we now turn to. Further down the section "Computational performance" is devoted to a closer look at the complexity and the time consumption.

##### 3.2 Unrestricted search

This more free approach to identifying particular geometries consists simply in looking for cases of rare writhe values. As explained in the main text, for a fixed sub-chain length we compute the mutual writhe of all pairs of such sub-chains; to avoid rather massive amounts of output, we pick out for each chain the case of lowest and the case of highest writhe value (the lowest being in general negative). With every high writhe value there will be several nearby sub-chains having almost the same high writhe, and to tackle this we here proceed a little brutally as just described. In the distribution of these extreme values over e.g. the top100 set we then consider the top-scoring cases. This implies of course that we will find at most two conspicuous examples per structure. While not the final version of such a search method, it should suffice for our purpose: to keep the search "open" while at the same time checking if we can re-discover the link examples we found in the restricted search.
Here follow results both for the top100 and the top8000 set. First top100 (Fig.S10):

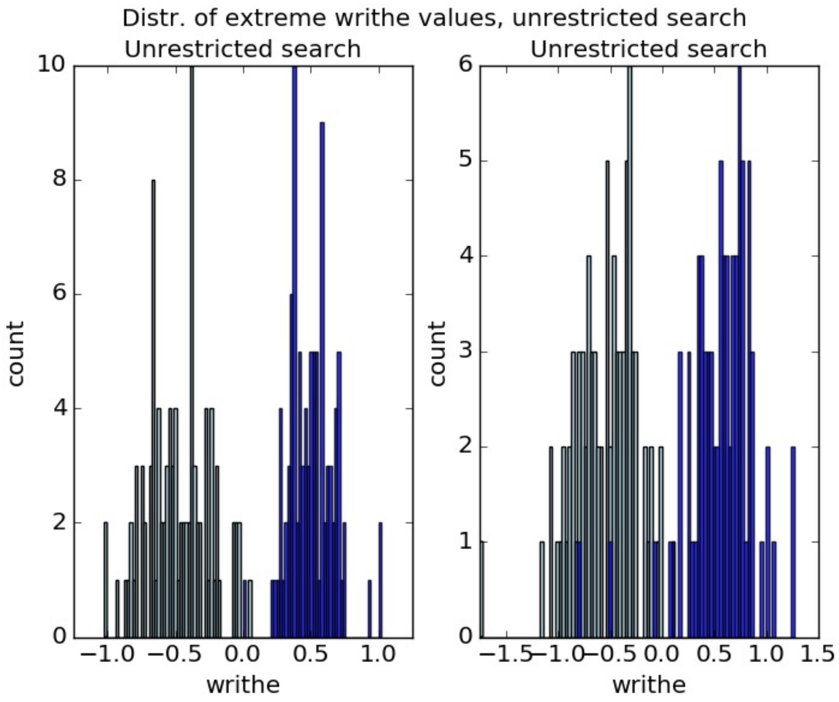


Fig.S10. Distributions of extreme writhe values for pairs of sub-chains; for each protein in the top100 set the highest (blue) and the lowest (possibly negative, lightblue) value are kept. For the plot to the left the fixed sub-chain length was set to 15; to the right the value was 30.

We notice from these plots that the range (almost) sits within $[-1, 1]$ for length 15, while for length 30 there are a few cases outside this interval and one rather extreme outlier below -1.5. Rather as a curiosity one may also observe that there are cases in which the maximal writhe value goes almost as low as -1. The reason for this is that in some very short structures there is only just enough room for two disjoint sub-chains of the set length and, coincidentally, in a couple of these the writhe of this pair is very low. Indeed, this phenomenon is seen for sub-chains of length 30, but not for length 15.

Let us go through the top-5 negative writhe value examples and then the top-5 positive, first for a sub-chain length of 15 and thereafter for length 30; we shall only show the examples that we have not already met in the restricted search above.

| Structure/chain | Pair | Mutual writhe | Type (by inspection) |
|---|---|---|---|
| 1dif/A | (23,38);(71, 86) | 1.02 | link |
| 1dif/B | (23,38);(71, 86) | 1.02 | link |
| 1kap/P | (51,66);(108, 123) | 0.93 | pseudo-link |
| 8abp/- | (222,237);(237, 252) | 0.76 | self-poke |
| 1lam/- | (370,385);(385, 400) | 0.75 | self-poke |
| 1arb/A | (152,167);(168, 183) | -1.02 | pseudo-link |
| 1bpi/- | (9, 24);(29, 44) | -1.02 | link |
| 7rsa/- | (71,86);(94,109) | -0.94 | pseudo-link |
| 1ptx/- | (0,15);(41, 56) | -0.86 | poke |
| 1lit/- | (92,107);(109, 124) | -0.85 | self-poke |

Table 4: Top 5 positive and top 5 negative writhe cases from the unrestricted search in the top100 based on sub-chains of lengths 15 (and implicitly step size 1). Pair refers to the indices of the segments in the chain bordering the two sub-chains.

The first example (Fig.S11a) for the length 15 case is one of a "sub-chain that winds on itself":

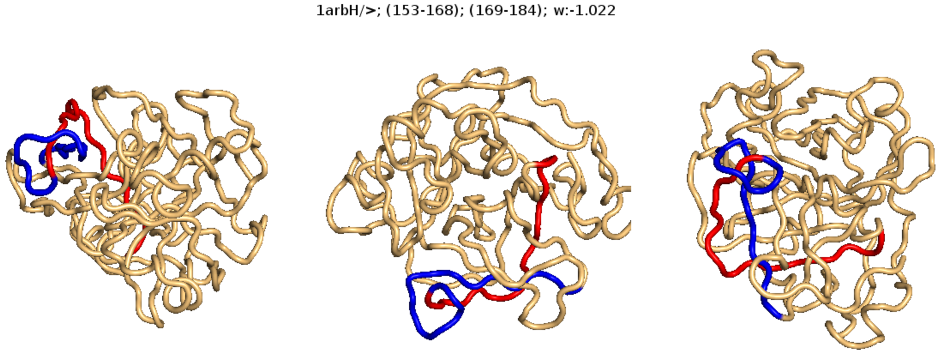


Fig.S11a. The geometry of the sub-chain pair in the 1arb protein of highest negative writhe (∼ -1.022) in the top100 set.

One may notice that, upon connecting the ends of the blue resp. the red strand by straight line-segments, this example becomes a 1-link, i.e. a "pseudo 1-link". The next example is the link in 1bpiH that we found as top-scoring in the restricted search. Then follows another example (Fig.S11b) of two sub-chains winding on one another (i.e. another pseudo 1-link):

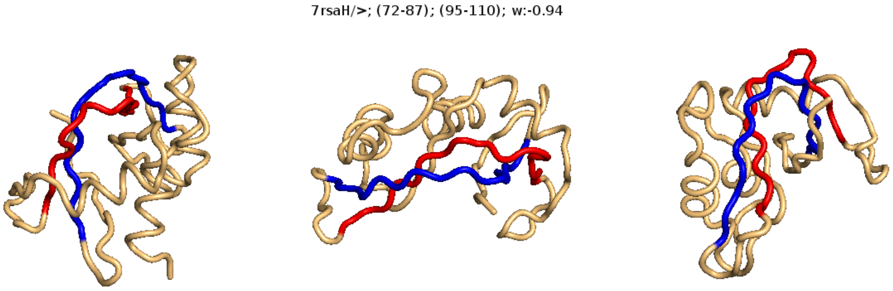


Fig.S11b. The geometry of the sub-chain pair of 3rd most negative writhe ($\sim$ -0.940) in the top100 set, found in the 7rsa protein.

The last two examples of the top-5 negative writhe values is a self-poke or pseudo-link (in 1lit of writhe about -0.85) and this (Fig.S11c):

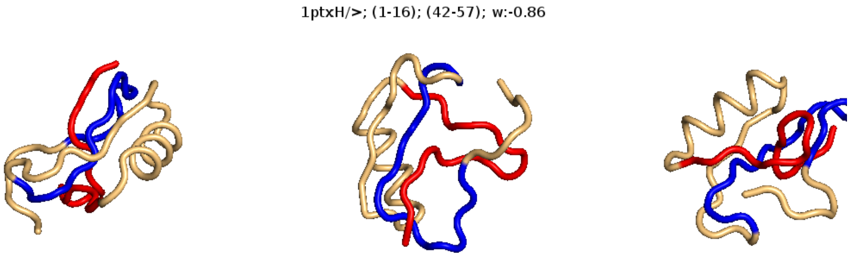


Fig.S11c. The geometry of pair of sub-chains of writhe $\sim$ -0.860 in the top100 set, found the 1ptx protein.

The top-5 positive values are headed by the highly similar 1-links in the A and B chains of 1dif (the writhe value is similar too: about 1.022 here and 0.968 above). Of these two of which the link in the A chain was found in the restricted search above, while the one in the B chain was not (the reason being that one of the two subchains does not qualify as almost closed). The following case is seemingly a pseudo-link:

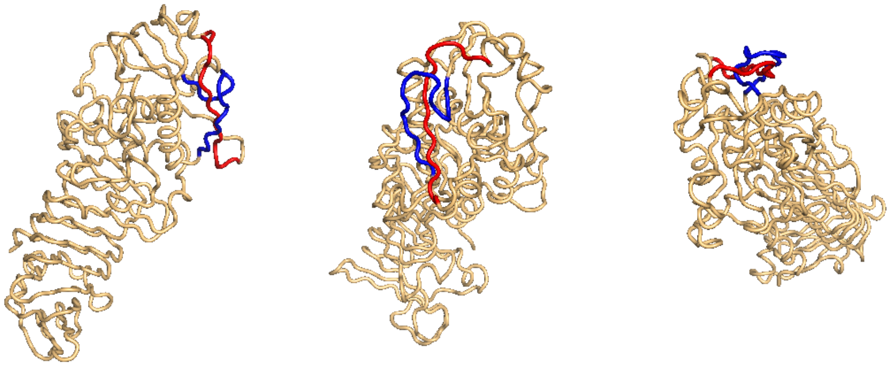


Fig.S11d. The geometry of the sub-chain pairs in the the 1kap protein making the 5th most positive writhe in ($\sim$ 0.929) the top100 set.

The 4th and 5th place are held by rather weak cases (low writhe) in the 8abp and 1lam structures.

So what emerges here is that we re-discover the cases found in the restricted search while adding a true link and then some pseudo 1-links which were not caught by the restricted method since the sub-chains do not qualify as closed loops.

When moving to the length-30 version the geometries sometimes become harder to decipher, but new interesting cases show up, too:

As above we start by the top-5 negative writhe value examples. Here the first (Fig.S12a) is a case of a "double poke"; two almost-loops aligning — or "sheeting" — each poking through the other (also shown in the main paper), while one subchain (colored blue in Fig.12a) winds around the other (colored red). While not possible here, turning this around for different views [9] can help revealing that this is not a case of a knot. Below, searching in the top8000 set with the GISA rar0 scan tool, a whole series of highly similar double-pokes are found, and of similar high writhe values too.

[9] By running the accompanying Python or Pymol scripts, plots allowing interactively rotating the structures can be had, or, if the highlighting of the subchains can be done without, by looking the structure up in PDB [5]

| Structure/chain | Pair | Mutual writhe | Probability |
|---|---|---|---|
| 1dif/B | (14,44);(65, 95) | 1.27 | link |
| 1dif/A | (14,44);(65, 95) | 1.26 | link |
| 1kap/P | (50,80);(102, 132) | 1.07 | pseudo-link |
| 2trx/A | (9,39);(39, 69) | 1.00 | self-poke |
| 2olb/A | (351,381);(381, 411) | 1.00 | self-poke |
| 2cpl/- | (70,100);(102, 132) | -1.75 | double-poke/pseudo-knot |
| 1nif/- | (230, 260);(260, 290) | -1.17 | self-poke |
| 1php/- | (239,269);(269,199) | -1.08 | self-poke |
| 2olb/A | (245,275);(464, 494) | -1.07 | poke |
| 1mla/- | (126,156);(162, 192) | -1.01 | self-poke |

Table 5: Top 5 positive and top 5 negative writhe cases from the unrestricted search in the top100 based on sub-chains of length 30 (and implicitly a step size of 1). Pair refers to the indices of the segments in the chain bordering the two sub-chains. Pair refers to the indices of the segments in the chain bordering the two sub-chains.

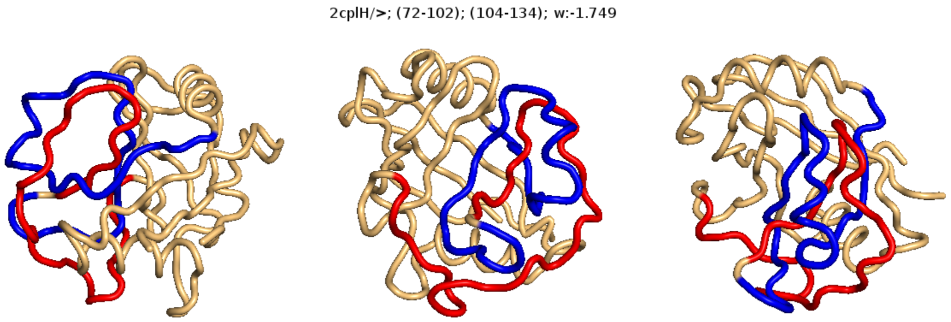


Fig.S12a. The geometry of the lenght-30 sub-chain pair in the 2cplH protein of highest negative writhe (∼ -1.749) in the top100 set.

The next (Fig.S12b) shows a loop followed by a sub-chain that aligns to the loop (i.e. another "sheeting"); the largest contribution to the writhe probably comes from the red poking through the blue (right "after" the red loop):

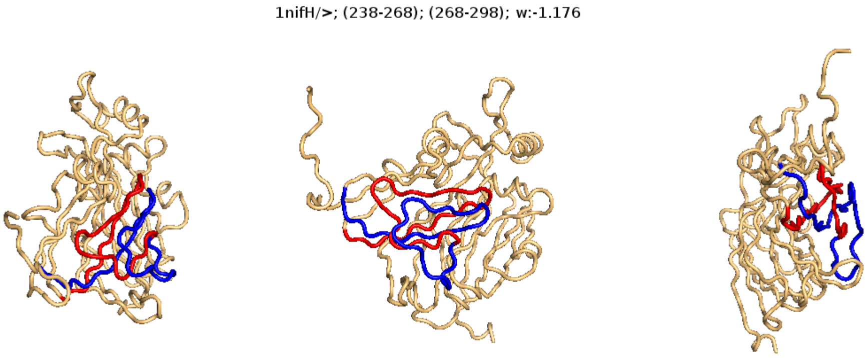


Fig.S12b. The geometry of the length-30 sub-chain pair of second highest negative writhe ($\sim$ -1.176) in the top100 set, found in the 1nif protein.

The following example appears to be a self-poke (in 1php), which we skip, then follows a straight poke in 2olb and then this (Fig.S12c), which could qualify as a 1-link while at the same time incorporating two short aligned helices:

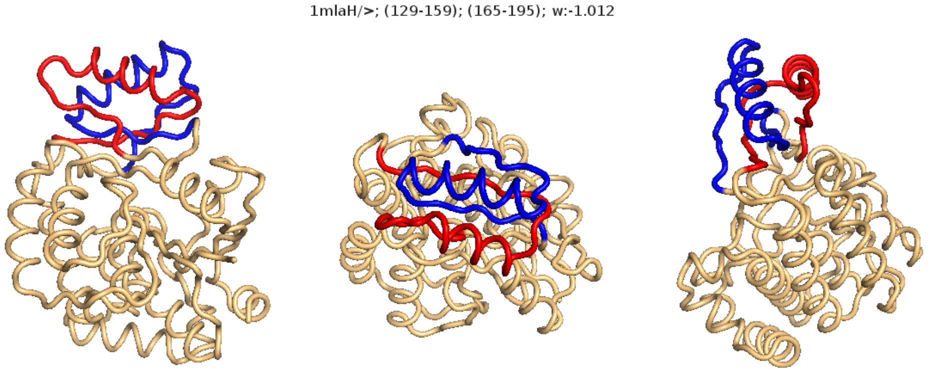


Fig.S12c. The geometry of the length-30 sub-chain pair in the 1mla protein of the 5th most negative writhe ($\sim$ -1.012) in the top100 set.

We may notice that the nice link in 1bpi was not found; this is can be ascribed simply to the fact that we are searching for disjoint sub-chains of length 30, for which there is then not enough room in such a small molecule (1bpi has a length of 58; as we saw above, the link in 1bpi was found using subchain length 15).

Turning now to the top-5 positive writhe value examples, the first are the 1-links in the A and B chain of 1dif, following these comes an extension (of writhe ∼ 1.07) of the poke in the 1kapH found with sub-chain length 15 (which has writhe ∼ 0.93). The final two cases are self-pokes in 2trx and 2olb.

To sum up on the unrestricted search in the top100 set, when moving to length-30 from length-15 (and from the restricted search in particular) we see that most cases are retained while new ones with more intricate geometry appear. So, as expected, the "heading or trailing pieces" of sub-chain strands that are added when moving to length-30 do not appear to be blurring the picture (as seen in these examples, e.g. in 1kap, the actual writhe values are not changed much by these additional pieces). Thus the advantage of an unconditional search seems to come true: without "prejudice" — we do not need to specify a geometry that we are looking for — particular shapes surface.

In the same vein, in the top8000 set which we now turn to, some more "interesting" geometries show up, in particular when searching with a sub-chain length of 30. With sub-chain length 15 we only see simple wind cases, the two sub-chains intertwining — a pseudo 1-link with maybe more than one winding (the case in 3ec0 could though be regarded rather as a straight poke). This single sided nature of the highest writhe cases (and of any sign) is somewhat remarkable. Here is an example:

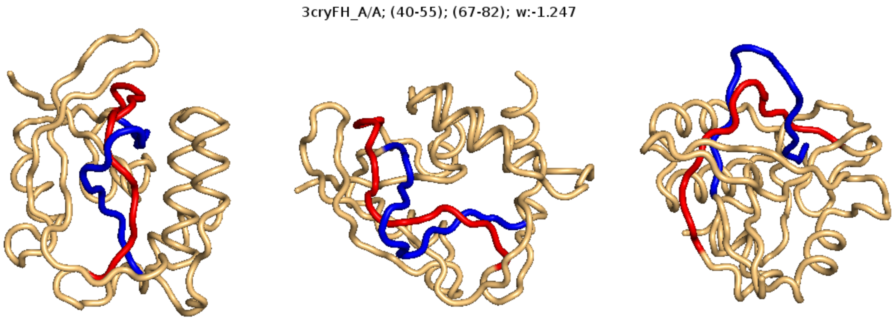


Fig.S13. The geometry of the length-15 sub-chain pair in chain A of the 3cryFH protein, of 5th most negative writhe (∼ -1.247) in the top8000 set.

Let us now focus on the search with a sub-chain length of 30 and consider the top-10 there (Table 6):

| Structure/chain | Pair | Mutual writhe | Type (by inspection) |
|---|---|---|---|
| 3onp/A | (60,90);(106,136) | 1.50 | knot |
| 2i6d/A | (164,194);(195,225) | 1.48 | knot |
| 1ual/A | (74,105);(105,135) | 1.47 | knot |
| 1ns5/B | (61,91);(92,122) | 1.47 | knot |
| 3o7b/A | (128,158);(161,191) | 1.44 | knot |
| 3hms/A | (0,30);(56,86) | -1.84 | double-poke |
| 2r99/A | (70,100);(102,132) | -1.76 | double-poke |
| 2cmt/A | (70,100);(102,132) | -1.76 | double-poke |
| 2wfj/A | (78,108);(110,140) | -1.76 | double-poke |
| 2cfe/A | (69,99);(101,131) | -1.76 | double-poke |

Table 6: Top 5 positive and top 5 negative writhe cases from the unrestricted search in the top8000 based on sub-chains of length 30 (and implicitly step size 1). Pair refers to the indices of the segments in the chain bordering the two sub-chains.

The geometries/topologies found here are

- Double-pokes, like the one in 2cpl above. Negative writhe only.
- True knots: the two sub-chains are adjacent and build a simple knot. Positive writhe only.

As we shall se below when considering the output from a rar0 scan in the top8000 set, these characteristics extend further down the top ranking (the top 10 positive writhe cases being knots, and by and large all in top 15 negatives being double-pokes and sharing the configuration).

Turning again to have a look examples, the highest negative writhe cases is this in 3hms, which appears to be something like a one-and-half wind case (a slip-knot, maybe):

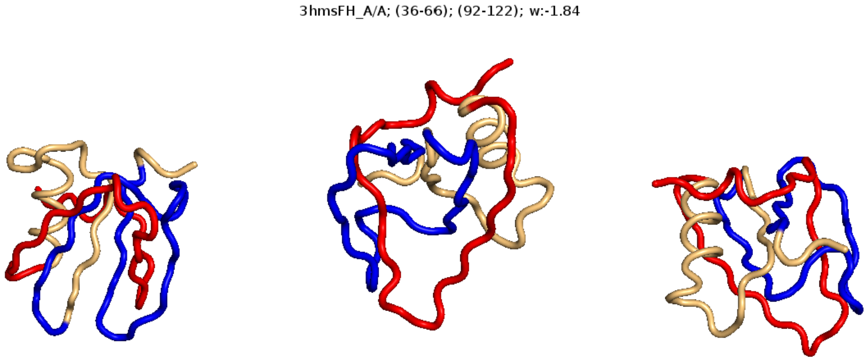


Fig.S14a. The geometry of the length-30 sub-chain pair in chain A of the 3hms protein, of highest negative writhe ($\sim$ -1.84) in the top8000 set.

The following four examples are slip-knots/double-pokes, like this (Fig.S14b) and similar to the one in 2cpl (Fig.S12a):

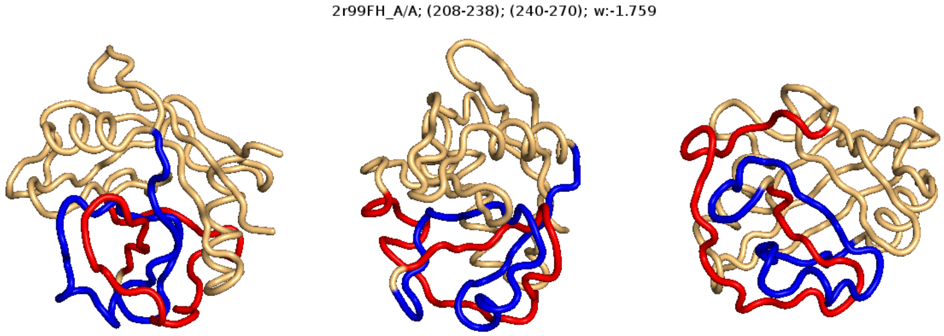


Fig.S14b. The geometry of the length-30 sub-chain pair in chain A of the 2r99FH protein, of 2nd most negative writhe ($\sim$ -1.759) in the top8000 set.

The positive writhe cases are all knots; here the first two (Fig.S14c-d):

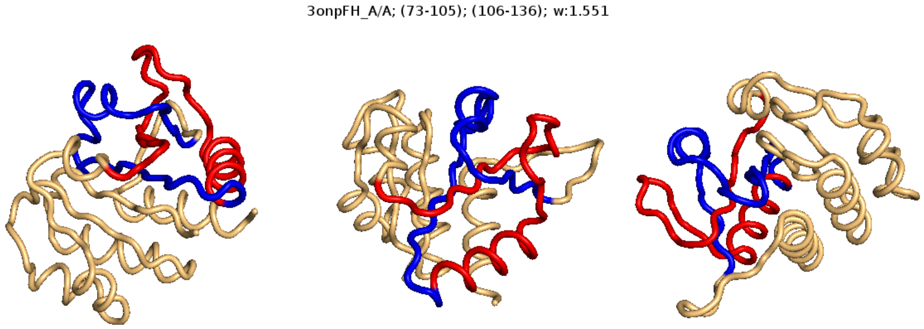


Fig.S14c. The geometry of the length-30 sub-chain pair in chain A of the 3onpFH protein, being of 2nd highest positive writhe ($\sim 1.551$) in the top8000 set.

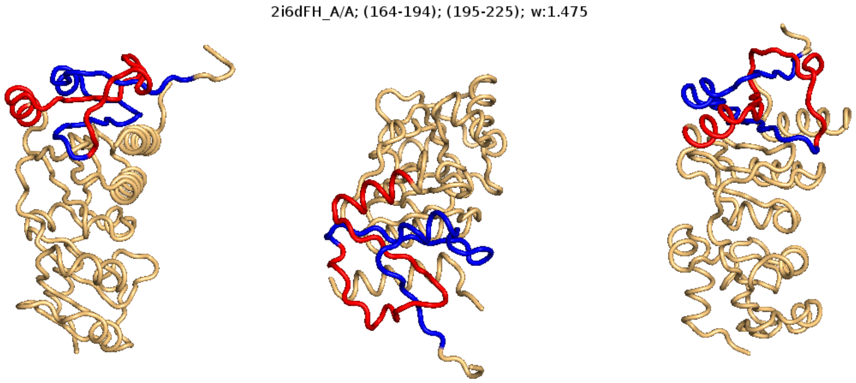


Fig.S14d. The geometry of the length-30 sub-chain pair in chain A of the 2i6dFH protein of 5th most positive writhe ($\sim 1.475$) in the top8000 set.

Do the links found in the restricted call still appear as rare in the unrestricted search, if at all present? Considering the 21 link cases having absolute mutual writhe above 0.95 (see text at Table 2), the answer is: yes, they are still present and with high mutual writhe values, but their rareness is somewhat shaken: of the 9 cases of positive writhe, six are placed in top

10 from the unrestricted search, 2qd6 is at rank 14, and only two are to be found further down the list, viz. 1pqh (writhe 14.01) and 3m3q (writhe 12.11). Of the 12 negative writhe cases, several are overlapping so only three are essentially different; of these 3hms is right at the top of the list from the unrestricted search, while 3dqp (writhe -14.92) and 3fdr (-12.63) are placed much further down. So the two latter link cases are drowning in the more elaborate configurations of higher negative writhe. Since the negative writhe cases here are so few, let us consider all 31 from the restricted search having absolute mutual writhe above 0.9. These make up some 9 additional cases, of which four appear with with very high writhe in the unrestricted search (2hq6, 2x7k, 2fu0, 2a2n), three have moderate writhe (1xlq, 1i7h, 2vve), one (2bt6) is of somewhat lower writhe (-8.80) and one (1taw) is too short to contain two disjoint subchains of length 30. As for the 2bt6 case, the link is made up by two (almost) adjacent subchains of length about 20, why two subchains of length 30 will cover it less well and the unrestricted search at length 30 therefore ascribes a lower writhe to it (and a run on a lower subchain length should also be made). Altogether it appears that the rareness is by and large retained for the positive writhe cases, while on the negative side many cases of higher negative writhe appear and "bury" the link cases.

##### 3.3 Results from GISA scans

We here focus on the basic scan method, rar0, of GISA, which amounts to a formalized and slightly extended version of the unrestricted search. We show a few results and include some checks of the two other methods, rar1 and rar2.

Let us first show the top 10 and the top 15 of the results from the rar0 scan of top8000 vs. itself, using only the cases of highest positive and the highest negative mutual writhe per structure (excerpts of these top rankings are shown in the main paper). We here expect to see the same cases as we found right above in the unrestricted search through top8000 and in the same order. That is indeed the case (Table 7).

According to the KnotProt-server [3] these 10 cases (Table 7) are all knots (we showed two examples above in Fig.S14c and d). The negative writhe cases (Table 8) are not identified by the KnotProt-server (which should probably also not be expected); all from rank 2 to 15 except the case in 2v25 are "double-pokes" and, by visual inspection, structurally very similar to the 2cpl case shown above (Fig.S12a) and in the main paper (see

| Structure/chain | Pair | Mutual writhe | Rank |
|---|---|---|---|
| 3onp/A | (60,90);(106,136) | 18.87 | 1 |
| 2i6d/A | (164,194);(196,226) | 18.49 | 2 |
| 1ual/A | (74,104);(104,134) | 18.32 | 23 |
| 1ns5/B | (62,92);(92,122) | 18.19 | 4 |
| 3o7b/A | (124,154);(162,192) | 18.08 | 5 |
| 2egv/A | (138,168);(178,208) | 17.62 | 6 |
| 2ha8/B | (58,88);(96,126) | 17.33 | 7 |
| 3aia/A | (112,142);(148,178) | 17.22 | 8 |
| 3n4j/A | (68,98);(98,128) | 17.08 | 9 |
| 2qmm/A | (104,134);(142,172) | 17.03 | 10 |

Table 7: Top 10 structures in rar0 ranking of the top8000 set vs top8000 as back ground, based on the highest positive mutual writhe pair per structure. Pair refers to the indices of the segments in the chain bordering the two sub-chains. Pair refers to the indices of the segments in the chain bordering the two sub-chains.

also Fig.S14b). These 14 proteins belong to a family of cis-trans isomerases. The shared configuration found here contains only little secondary structure (cf. the figures).

As is apparent from Table 8 these 14 cases (configurations) sit in very similar places in the structures. We then made a multiple alignment of the 14 structures, using ClustalOmega 2.1 [9] for the purpose. In the resulting alignment, both the complete sequences and the considered sub-sequences appeared well aligned (i.e. the sub-sequence containing the two sub-chains of the configuration). Of the about 60 residues in the sub-sequence, 20 were fully conserved and some further 10 showed good but not perfect similarity. So, overall, the sequence similarity is roughly 50 pct., which though does not appear high considering the similarity of the folds in this region, which is even of low secondary structure content.

To make a comparison with the unrestricted search in the top100 set (Table 5) , we list (Table 9) the top ranking structures from a rar0 scan of the top100 set against top8000 as background.

| Structure/chain | Pair | Mutual writhe | Rank |
|---|---|---|---|
| 3hms/A | (0,30);(56,86) | -23.13 | 1 |
| 2r99/A | (70,100);(102,132) | -22.10 | 2 |
| 2cmt/A | (70,100);(102,132) | -22.09 | 3 |
| 2wfj/A | (78,108);(110,140) | -22.08 | 4 |
| 2igv/A | (78,108);(110,140) | -22.04 | 5 |
| 2esl/A | (74,104);(106,136) | -22.01 | 6 |
| 2z6w/A | (70,100);(102,132) | -22.01 | 7 |
| 3k2c/B | (70,100);(102,132) | -21.92 | 8 |
| 2v25/A | (70,100);(162,192) | -21.65 | 9 |
| 1xo7/B | (72,102);(104,134) | -21.64 | 10 |
| 2a2n/C | (70,100);(102,132) | -21.52 | 11 |
| 2hq6/A | (70,100);(102,132) | -21.47 | 12 |
| 2cfe/A | (70,100);(102,132) | -21.26 | 13 |
| 1zkc/A | (70,100);(102,132) | -21.16 | 14 |
| 3ich/A | (76,106);(108,138) | -21.12 | 15 |

Table 8: Top 15 structures in rar0 ranking of the top8000 set vs top8000 as back ground, based on the highest negative mutual writhe pair per structure. The sub-chain length was 30 and the step size 2. Pair refers to the indices of the segments in the chain bordering the two sub-chains.

The rankings here (Table 9) are almost as from the unrestricted search (Table 5) ; the top 3 are the same in the two lists, while the following e.g. four are shared but come in different ordering (for the postive writhe 2ctc, 2olb, 2trx and 2tca are placed next, and for the negatives 1mla, 2olb, 1rcf, 1tta); the writhe values for these cases are quite similar, so the reason for these different orderings simple come from the different step sizes used (in the unrestricted search the step size is 1, while we used a step size of 2 in the rar0 run).

Let us finally consider the rar1 and rar2 scans briefly. To check the sanity of the methods we ran a few tests:

- Running rar1 (rar2) normalized vs. unnormalized, using only the writhe with no threshold (done with a negative threshold)
- Comparing the output from rar1 (rar2) against that of rar0, using only the writhe, unnormalized.

| **Structure/chain** | **Pair** | bfMutual writhe | **Probability** |
|---|---|---|---|
| 1dif/B | (12,42);(64, 94) | 15.77 | $2.310^{-3}$ |
| 1dif/A | (12,42);(64, 94) | 15.70 | $2.510^{-3}$ |
| 1kap/P | (50,80);(102, 132) | 13.48 | $8.410^{-3}$ |
| 2ctc/A | (186,216);(244, 274) | 11.33 | $4.710^{-2}$ |
| 2olb/A | (352,382);(382, 412) | 10.99 | $6.510^{-3}$ |
| 2cpl/- | (70,100);(102, 132) | -21.98 | $1.010^{-3}$ |
| 1nif/- | (230, 260);(260, 290) | -14.77 | $1.810^{-2}$ |
| 1php/- | (240,270);(270,300) | -13.09 | $4.210^{-2}$ |
| 1mla/- | (126,156);(162, 192) | -12.71 | $4.910^{-2}$ |
| 2olb/A | (244,274);(464, 49 4) | -12.57 | $5.310^{-2}$ |

Table 9: Top 5 ranking positive and top 5 ranking negative writhe cases from a rar0 scan of the top100 set against top8000 as back ground using sub-chains of length 30 and a stepsize of 2. Pair refers to the indices of the segments in the chain bordering the two sub-chains.

The first of these showed no differences between the normalized and unnormalized results. For the second we here show scatter plots (Fig.S15) of the scores from the three scans, used at comparative settings:

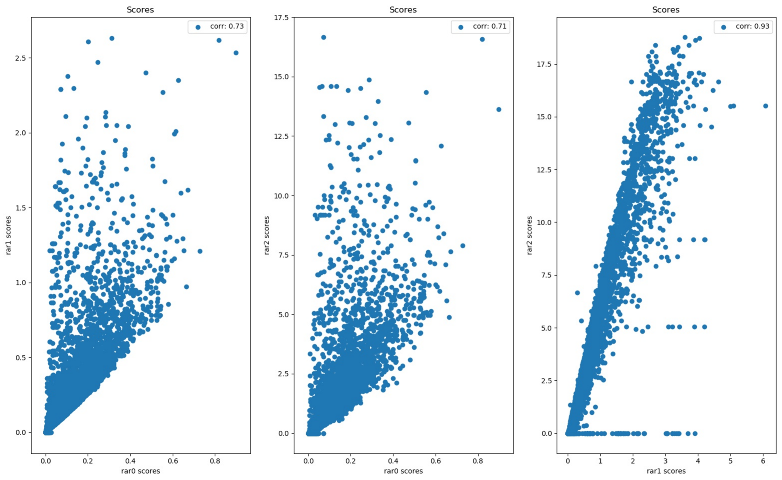


Fig.S15. Comparisons of the scores output by rar0, rar1 and rar2 when run on the top8000 set against itself. For the description of each plot see the text.

In the two left-most plots (of Fig.S15) rar0 is compared to rar1 and rar2 respectively, using only the writhe; in the right-most plot rar1 is compared to rar2 using the writhe and average crossing number for rar1 and the pairs matching in rar2, and the first five invariants for the single window matching in rar2. No mismatches were allowed in any of the runs.

Evidently, the scores correlate well. The expectation here is that the scores from rar1 will correlate better with those from rar0 than will those from rar2 (since rar1 operates directly on the pairs, while the scoring in rar2 goes via the single window matches) and that scores from rar1 and rar2 will be better correlated than those from rar0 (as their scoring methods are more similar than to that of rar0). These overall tendencies are seen.

##### 3.4 Computational performance

The recursion formulas clearly suggest that the computational complexity of the base algorithm for computing the GIs of order less than three should be $O(L^2)$, while $O(L^3)$ in order three (with $L$ the length of the chain). To show that this is indeed the case we have collected the average computation time per protein chain in the top100 and in the top8000 set over 100 and over 20 repeated runs, respectively. The time consumption covers the computation parts without invoking the search code (i.e. closed_loops_b = 0 and invValSubChainPairs_b = 0), and excludes the initial load of the structures and accompanying allocation of memory. The time estimates below were obtained running a Windows-runnable version of the code on a common laptop (Intel Core i7-4510, 2.00 GHz/2.60GHz, 8GB RAM, hard disc of SSD type; OS Microsoft Windows 10).

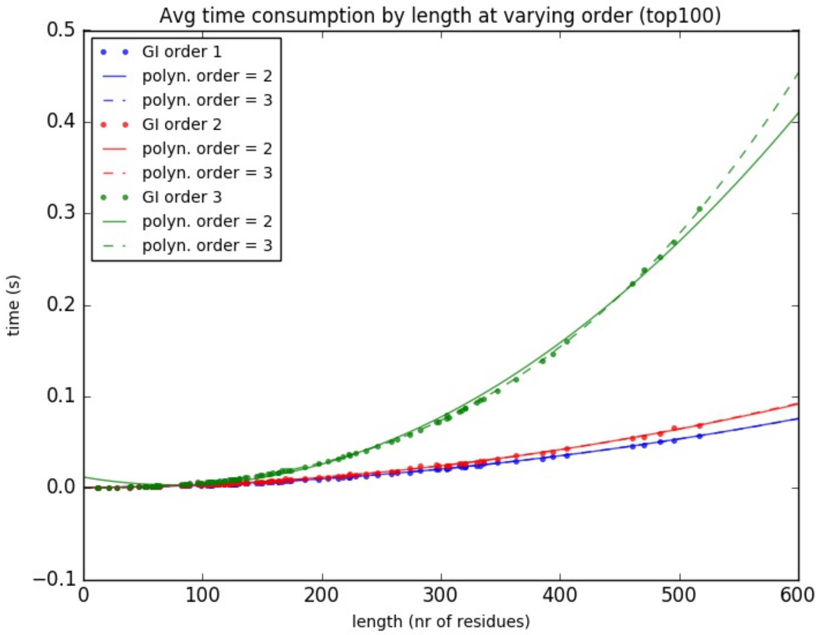


Fig.S16a. Average computation time per chain over 100 repeated runs on the top100 set. The computation returns invariant values at the indicated order (GI order) across the full simplex (see more in the text). Dots indicate observed data, while the dashed and full lines show best fit polynomiums of the indicated order (obtained with Python Numpy.polyfit). Colors follow the GI order.

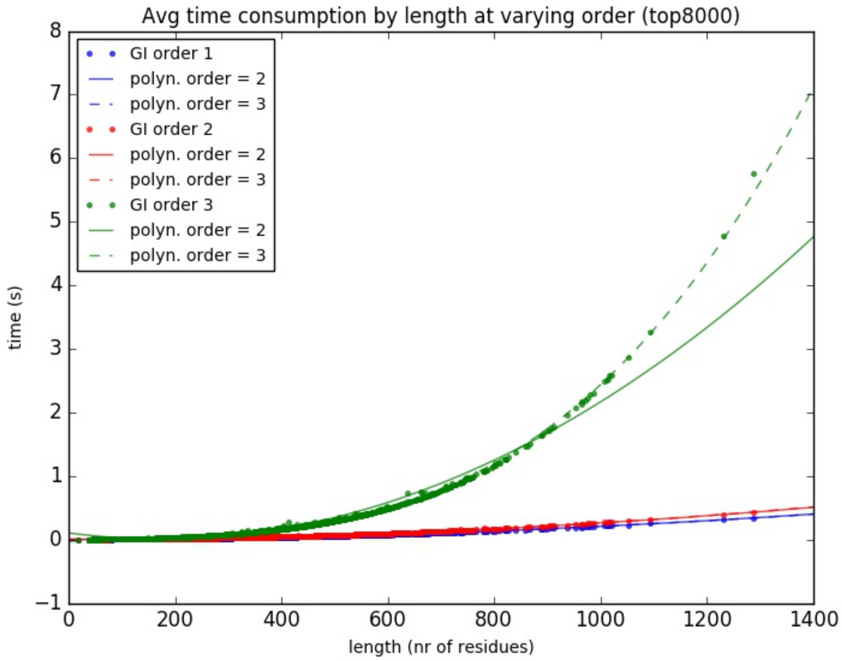


Fig.S16b. As Fig.S16a but for 20 repeated runs on the top8000 set.

Clearly, for the GIs of order one and two a 2nd order polynomium gives a good fit, while in order three the best fit is obtained with a 3rd order polynomium (Fig.S16a-b).

This complexity implies that getting the additional local GI values does not result in a severe time consumption. Thus, in test runs the performance of GISA compared well with that of the algorithm in [7] (which in order 2 and 3 only computes the GIs on the global structures). It is in its place here to elobarate on the output of GISA. When run in (GI) order one (i.e. with the parameter "order" set to 1), GISA produces the order one GIs on the full chain as well as on all connected sub-chains. When run in (GI) order two, GISA produces in addition the GIs of order two on the full chain along with some new 2nd order invariants of relative type on all sub-chains. When run in (GI) order two "full" mode (i.e. with the parameter "full_b" set to 1), GISA produces in addition the GIs of order two on all sub-chains. Finally, when run in order 3, GISA's output contains in addition the order three GIs on the full chain along with some new 3rd order invariants of relative type on all connected sub-chains. The computational complexity in order two full mode is the same as in (GI) order three, i.e. $O(L^3)$.

To examine the additional time spend on computations done for the searches we ran repeats as above, but now with the search code invoked. The time consumption shown is the expected (i.e. average) search time including the time spend on computing the GIs, e.g., when the search is performed on one out of many protein models and the one model is presently held in memory. As writing the results to a file can be costly, we put write-out thresholds so as to limit the print out to a few cases, if any. Clearly, the additional time consumption is small (Fig.S17a-b):

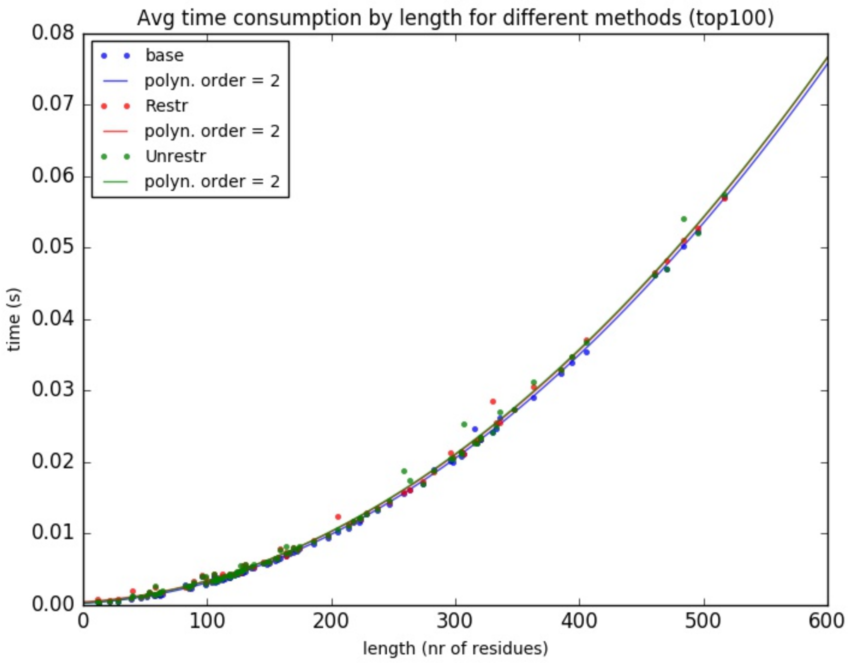


Fig.S17a. Average computation time per chain over 100 repeated runs on the top100 set when invoking the searches or not (base). The computation returns the GIs of order one across the full simplex (i.e. on all connected sub-chains). Dots indicate observed data, while full lines show best fit polynomiums of the indicated order (obtained with Python Numpy.polyfit). Colors follow the search method (none being the base case).

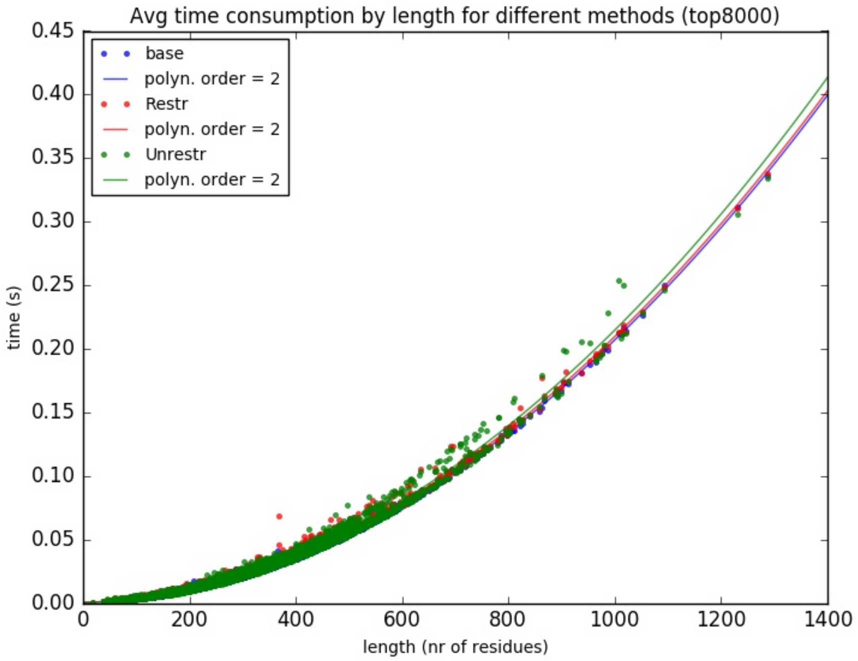


Fig.S17b. As Fig.S17a but for 20 repeated runs on the top8000.

Roughly from these plots, an unrestricted search on top of a computation of the GIs in order one adds less than 5 pct to the time consumption. As expected the similar overhead is slightly lower for the restricted search. In order two and three (not shown) this constant overhead is then even lower compared to the base computation time.

To support this, the time for completing the 100 repeats of the base computation on the top100 set was 89 s, 101 s and 352 s, for GI order one, two and three respectively. With the search code invoked (either one) the similar numbers were 93 s, 105 s and 360 s. On the top8000 set the 20 repeats of the base computation took about 2214 s, 2542 s and 10993 s, for GI order one, two and three respectively. With an unrestricted search the same numbers were about 2344 s, 2681 s and 11059 s. (That the numbers do not simply scale by a factor of about 16 between the two data sets, can be ascribed to differences in the length distributions of the two sets; for instance the average length in the top8000 set is about 235 residues compared to 185 in the top100 set.)

### References


[1] F.Khatib and C.A.Roll and K.Karplus (2009). Pokefind: a novel topological filter for use with protein structure prediction. *Bioinformatics*, **25**, 281–288.

[2] Kinemage (2016). `kinemage.biochem.duke.edu`.

[3] KnotProt (2019). `https://knotprot.cent.uw.edu.pl/`.

[4] P. Røgen and B.Fain (2003). Automatic classification of protein structure by using gauss integrals. *PNAS*, **100**, 119–124.

[5] PDB (2016). `http://www.rcsb.org`.

[6] P.Røgen and H.Bohr (2003). A new family of global protein shape descriptors. *Math. Biosciences*, **182**, 167–181.

[7] Røgen, P. (2005). Evaluating protein structure descriptors and tuning Gauss integral based descriptors. *J. Phys.: Condens. Matter*, **17**, S1523–S1538.

[8] Røgen, P. *et al.* (2003). A new family of global protein shape descriptors. *Math. Biosciences*, **182**, 167–181.

[9] Sievers, F. *et al.* (2011). Fast, scalable generation of high-quality protein multiple sequence alignments using Clustal Omega. *Molecular Systems Biology*, **7:539**.
